# Alternative splicing decouples local from global PRC2 activity

**DOI:** 10.1101/2023.04.30.538846

**Authors:** Niccolò Arecco, Ivano Mocavini, Enrique Blanco, Cecilia Ballaré, Sophie Bonnal, Manuel Irimia, Luciano Di Croce

## Abstract

The Polycomb repressive complex 2 (PRC2) mediates epigenetic maintenance of gene silencing in eukaryotes via methylation of histone H3 at lysine 27 (H3K27). This complex interacts with sets of accessory factors to form two distinct subtypes, PRC2.1 and PRC2.2, that differ in their actions and chromatin targeting mechanisms. The underlying molecular mechanisms orchestrating PRC2 assembly are not yet fully understood. Here, we report that alternative splicing (AS) of the PRC2 core component SUZ12 generates an uncharacterised isoform SUZ12-S, which co-exists with the canonical SUZ12-L isoform in the same cell in virtually all tissues and developmental stages. We show that SUZ12-S drives PRC2.1 subtype formation and favours PRC2 dimerisation. While SUZ12-S is necessary and sufficient to ensure correct repression of Polycomb target genes via promoter-proximal H3K27me3 deposition, SUZ12-L is needed for maintaining global H3K27 methylation levels. Lastly, we show that mouse embryonic stem cells (ESCs) lacking either of the two isoforms have a slower exit from pluripotency upon differentiation stimulus. Our findings thus reveal a physiological mechanism regulating PRC2 assembly and higher-order interactions, with impacts on H3K27 methylation and gene repression.

## Introduction

PRC2 is responsible for the mono-, di- and tri-methylation of histone H3 lysine 27 (H3K27me1/2/3). This post-translational modification (PTM) helps to maintain gene silencing in a plethora of developmental processes, including pattern specification, X-chromosome dosage compensation and non-canonical genetic imprinting ^1^. The PRC2 core comprises four proteins: EZH1/2, EED, SUZ12 and RBBP4/7. The EZH2 SET domain is responsible for PRC2 histone methyl-transferase activity, although the formation of a minimal core complex including EED and SUZ12 is necessary for a fully-functional active site ^2–4^. The core complex is recruited to specific loci thanks to various accessory factors, including PCL1–3, which mediate the recruitment to unmethylated CpG islands (CGIs) ^5^, and AEBP2 and JARID2, which bind the PRC1-deposited mark of ubiquitinated H2A ^6,7^. Accessory proteins interact with PRC2 in sets, whereby the PRC2.1 subtype contains EPOP or PALI1/2 and one of the PCL family members, while the PRC2.2 subtype contains AEBP2 and JARID2^8,9^. These two subtypes co-localise at the vast majority of chromatin targets in ESCs ^10^. Disruption of the balance between the PRC2.1 and PRC2.2 subtypes leads to aberrant H3K27me3 deposition and transcriptional deregulation of target genes ^11,12^, suggesting the existence of PRC2 sub-type–specific functions. In addition to its specific composition, the ability of PRC2 to dimerise may also play a role in shaping its activity on chromatin ^13–15^. Alternative splicing (AS) allows eukaryotic cells to functionally diversify their proteome ^16^. We previously reported the existence of a class of alternative exons (termed ‘PanAS’) that differ from canonical AS switches in that they undergo AS in a non-tissue, non-developmental stage specific manner, producing broadly expressed alternative isoforms that co-exist in the same cell ^17^. Of note, these exons are in genes strongly enriched for chromatin-related functions, including various DNA-binding transcription factors (TF) and key histone-modifying complexes, such as PRC2. Here, we investigated the potential impact of PanAS events on the biology of PRC2. Among several events of interest, we focused on a novel, eutherian-specific SUZ12 isoform generated by PanAS skipping of exon 4. We demonstrate that this shorter isoform promotes PRC2.1 formation as well as dimerisation. Mechanistically, this isoform is responsible for the highly focused H3K27me3 deposition that is necessary to maintain Polycomb-mediated gene repression in ESCs.

## Results

### *SUZ12* contains an eutherian-specific PanAS exon

Inspection of the AS database VastDB ^17^ revealed various alternatively spliced exons among human PRC2 components with PanAS characteristics (e.g., widespread co-existence of alternative isoforms with no tissue-specificity or developmental stage–specificity) (Figure 1A; Table S1; see Methods). Some of these exons are predicted to disrupt the open reading frame when included or excluded, likely resulting in the occurrence of a premature stop codon. Others potentially generate functional alternative isoforms, such as the previously described AS events for the exons 4 and 14 of *EZH2* ^19–21^. Among these exons, AS of SUZ12 exon 4 is predicted to generate two alternative isoforms differing by the inclusion or exclusion of 69 nucleotides, which we named *SUZ12-long* (*SUZ12-L*) and *SUZ12-short* (*SUZ12-S*) respectively (Figure 1B). The 3D structure of PRC2 highlights the central role of SUZ12 as a scaffold for the assembly of the core complex and PRC2 subtype specification ^7,18,22^. In addition, SUZ12 mediates PRC2 dimerisation ^14,15^. Given its potential implications for PRC2 composition and function, we further characterised this AS event. The percentage of sequence inclusion (PSI) of the *SUZ12* exon 4 ranges mostly from 60% to 95% (median PSI, 86%) in both human and mouse across all adult and embryonic tissues (Figures 1C, 1D, and S1A), and analysis of full-length single-cell RNA-seq data from early development stages in human ^23^ or mouse ^24^ confirmed that both isoforms co-exist in the same cell (Figure S1B). In contrast, *suz12a* exon 4 is not alternatively spliced in zebrafish (median PSI, 100%), according to VastDB and RT-PCR assays using independent RNA samples (Figures 1C and 1D). Indeed, although exon 4 is highly conserved in length and sequence in all jawed vertebrates (Figures S1C and S1D), comparative analysis of the inclusion patterns across numerous vertebrate species revealed that this exon is alternatively spliced only in eutheria (placental mammals), while it displays full inclusion in all other species (Figure 1E).

**Figure 1.**
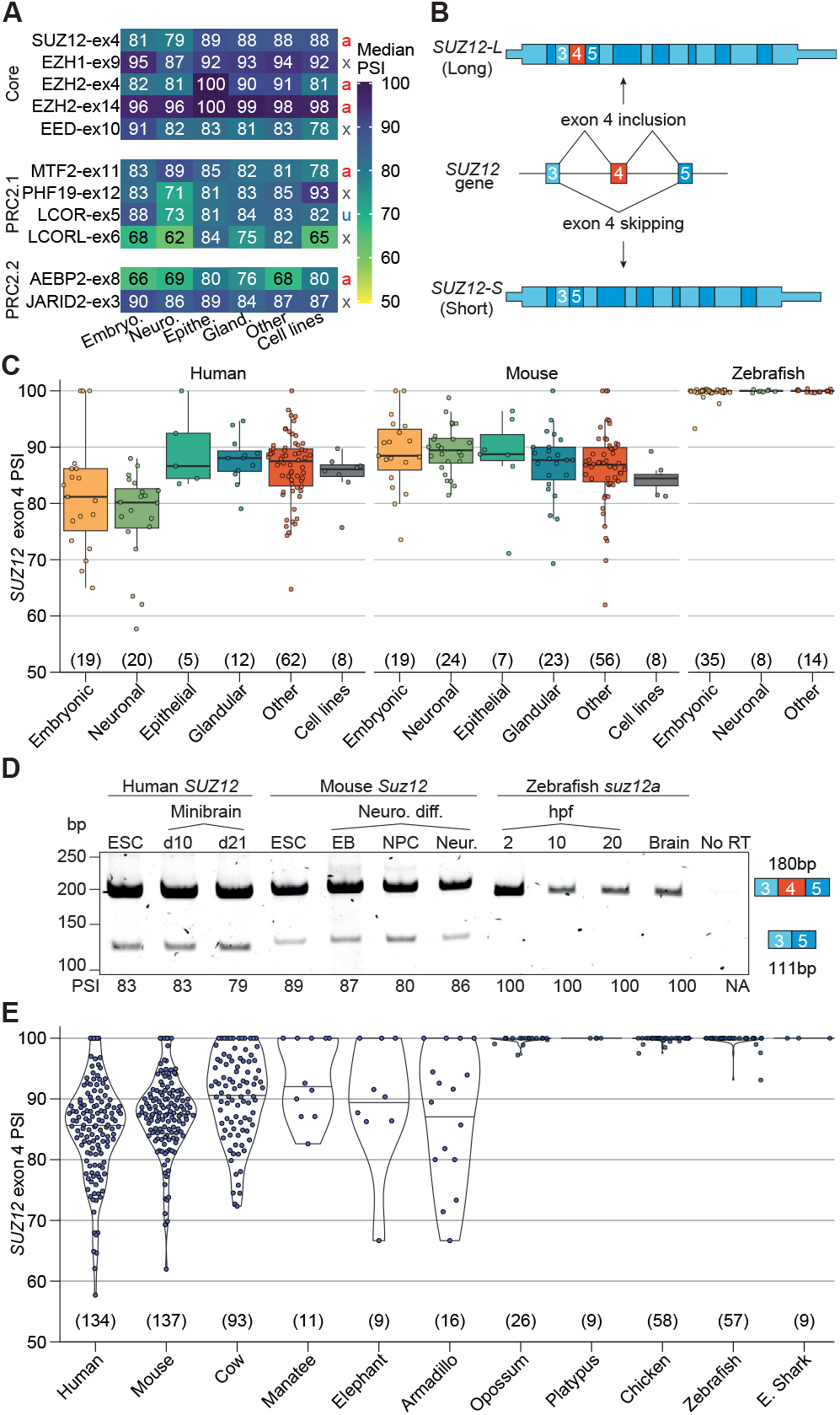
The PRC2 core component SUZ12 has an eutherian-specific alternatively spliced exon. (A) Heatmap showing median percentage of sequence inclusion (PSI) of AS exons in PRC2 components across different human tissues and cell lines. a, alternative isoform; x, ORF disrupted upon exclusion; u, UTR. (B) Schematic representation of two SUZ12 transcript isoforms generated by exon 4 inclusion (top) or skipping (bottom) from the SUZ12 gene (centre). GenBank IDs; SUZ12-L, NM_015355.4; SUZ12-S, NM_001321207.2. (C) Quantification of the SUZ12 exon 4 PSI across different tissue-types in three species. (D) RT-PCR of the SUZ12 exon 4 in human, mouse and zebrafish across neurodevelopmental timepoints. PSI is shown at the bottom, and the expected band size in human, at the right. (E) SUZ12 exon 4 PSI quantification across 11 vertebrates. The number of samples analysed is shown in brackets. All data points are presented in (C) and (E). The upper, centre and lower lines of the boxplot indicate the 75%, 50%, and 25% quantiles, respectively. Whiskers extend to the most extreme data point within 1.5× of the interquartile range.

### *Suz12* exon 4 is necessary for PRC2.2 formation

*Suz12* exon 4 encodes 23 amino acids (aa 129–152 in SUZ12-L) that partially overlap with the WD-binding domain 1 (WDB1, 110–145) (Figure 2A). Together with WDB2, this domain is responsible for accommodating RBBP4/7 into the lower lobe of PRC2^22^. In addition, this peptide provides a flexible hinge connecting the two SUZ12 domains that are essential for its association with accessory factors: the ZnB domain (aa 79–106) and the C2 domain (aa 153–365). The ZnB domain, together with the ZnF (aa 426–492), provide the binding surface for the EPOP C-terminal region (CTR) or the JARID2 transrepression (TR) domain. The C2 domain interacts with either PCL1–3 chromodomain or with the AEBP2 C2-binding (C2B) domain (Figure 2B). Therefore, we reasoned that exon 4 skipping might alter the structure of SUZ12 and, possibly, PRC2 composition. To explore this possibility, we generated ESCs that lack the *Suz12* exon 4 via CRISPR-Cas9–induced deletion (Figure S2A). We isolated three clones with the deletion (Δex4 #1–3) and two clones with no exon deletion (WT #1, 2; Figures S2B and S2C). We confirmed that Δex4 #1–3 clones exclusively expressed the short *Suz12* isoform at the mRNA and protein levels but maintained expression levels of the gene similar to the WT cells (Figures 2C and S2D). The presence of both protein isoforms in WT cells, and the absence of SUZ12-L exclusively in Δex4 cells, was further confirmed by precise reaction monitoring (PRM) targeted mass spectrometry (MS) (Figures 2D and S2E). Next, we performed SUZ12 immunoprecipitation coupled with mass spectrometry (IP-MS) in the WT and Δex4 clones to compare their interactomes. As expected, no peptides corresponding to exon 4 were retrieved in Δex4 samples, while the rest of the sequence displayed similar coverage (Figure S2F). Comparison of interactors in the two conditions revealed that SUZ12 binding to AEBP2 and JARID2 was strongly reduced in Δex4 cells with respect to WT cells, whereas SUZ12 binding to most core components or to PRC2.1-specific factors was unchanged or only slightly increased (Figures 2E and 2F; Table S3). These observations were confirmed by SUZ12 IP followed by Western blot (WB) (Figure 2G). Of note, the levels of the AEBP2 protein abundance were greatly reduced upon exon 4 deletion, likely due to its detachment from chromatin upon exclusion from the PRC2 complex (Figures 2G and S2G). To rule out any biases due to the expression levels and/or SUZ12 epitope masking, we generated Suz12 knockout (KO) ESCs (herein, KO) in which *Suz12* expression was subsequently rescued by re-introducing either the *Suz12-L* or *Suz12-S* mouse isoform fused to a triple-Flag tag under the regulation of a CAG promoter (KO+L/S; Figures S2H–S2K). Flag IP-MS in these cells confirmed that, while the long isoform was able to correctly form comparable amounts of both PRC2.1 and PRC2.2 subtypes, interaction of the SUZ12-S with PRC2.2-specific factors was drastically reduced (Figures S2L and S2M; Table S3). Overall these results indicate that only SUZ12-L is able to properly assemble both PRC2 subtypes, whereas the SUZ12-S-containing PRC2 preferentially assembles the PRC2.1 subtype.

**Figure 2.**
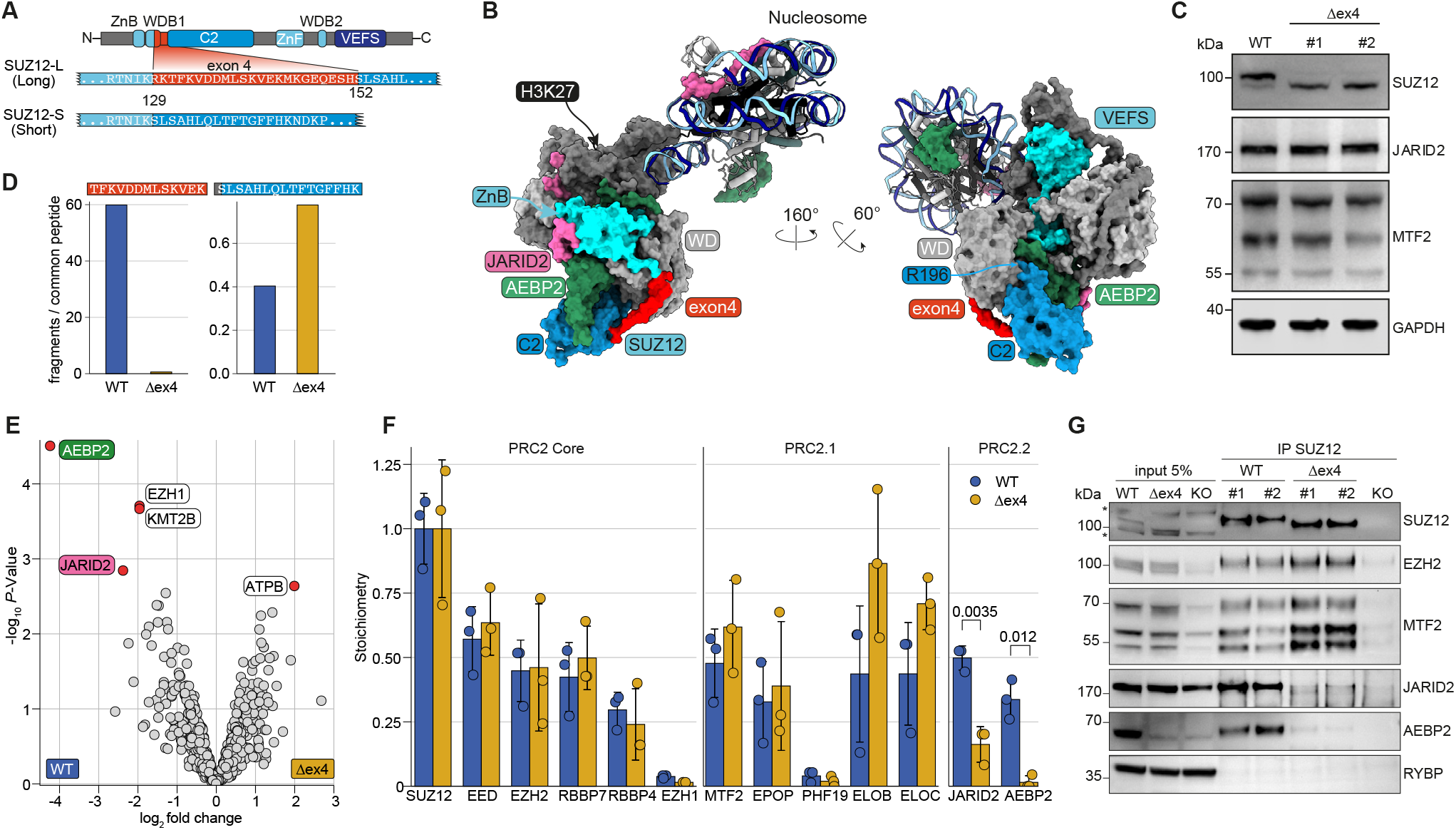
*Suz12* exon 4 skipping promotes PRC2.1 formation and dimerisation. (A) Schematic depiction of SUZ12 protein highlighting the relative position and amino acid (aa) sequence encoded by exon 4. Gen-bank IDs SUZ12-L: NP_056170.2; SUZ12-S: NP_001308136.1. (B) PRC2.2 structure surface rendering engaging a nucleosome according to ^18^. Region encoded by exon 4 is shown in red, the SUZ12 domains C2, ZnB, and VEFS and SUZ12 Arg 196 in cyan, histone H3 Lys 27, in black and the RBBP4 WD propeller, in grey. (C) WB showing SUZ12 protein abundance in WT#2 and Δex4 cells. Δex4 samples display only the lower band corresponding to SUZ12S. (D) Bar plot quantifying the SUZ12-L specific and SUZ12-S specific tryptic peptides acquired with PRM targeted proteomics. (E) Volcano plot of SUZ12 IP-MS in Δex4 vs WT ESCs. Significant proteins (adj. *P*-value *≤* 0.05, log_2_ FC > |1.5|) are shown as red points. Data were analysed using empirical Bayes statistics on protein-wise linear models using limma in DEP (see Methods). AEBP2 and JARID2 coloured as in (B). (F), Stoichiometric ratio of PRC2 core, PRC2.1 and PRC2.2 interactors between WT and Δex4 cells relative to bait (SUZ12). Points indicate individual biological replicates (n = 3 clonal cell lines); error bars: sd. Data were analysed using PSMs from *proteomicslfq*. Statistics: t-test. (G), IP-WB of SUZ12 in WT, Δex4 or KO ESCs.

### PRC2 dimerisation is favoured by SUZ12-S

PRC2 can form dimers ^13–15^ which are likely mediated primarily by a domain-swapping interaction that involves SUZ12 R196 (contained in the C2 domain) binding to the WD-repeat in the RBBP4/7 acidic pocket. Comparing the SUZ12 structures in the monomeric and dimeric contexts suggests that the flexibility of the C2 domain is likely influenced by the length of the hinge encoded by exon 4 (Figure S3A). Hence, we reasoned that the inclusion or exclusion of SUZ12 exon 4 may confer different dimerisation propensities to the complex. To test this hypothesis, we fractionated protein complexes from either WT or Δex4 nuclear extracts on a glycerol gradient (Figures 3A, 3B, S3B, and S3C). In WT extracts, we were able to distinguish EPOP- and JARID2-enriched PRC2-containing fractions, likely corresponding to PRC2.1 and PRC2.2 subtypes (fractions 10–12 and 12–16 respectively). Instead, in Δex4 nuclear extracts, we observed loss of JARID2 from the PRC2-containing fractions (fractions 6–8, Figures 3A, 3B, S3B, and S3C) in accordance with the IP-MS data. Interestingly, the bands corresponding to SUZ12-S, EZH2 or EPOP displayed a shift towards higher molecular weight fractions (14–18), compatible with a doubling in complex size (Figures 3B, 3C, and S3C). Similarly, isoform-specific rescue cell lines showed that the SUZ12-S assembles a higher-order PRC2 complex with respect to SUZ12-L (Figures 3D). Importantly, fractionation of a SUZ12 mutant unable to dimerise (SUZ12^3D^, K195D-R196D-K197D) ^14^ was comparable in size to the one of our long isoform (Figures 3D, 3E, and S3D). Altogether, these data suggest that *Suz12* exon 4 splicing influences the ability of PRC2 to dimerise.

**Figure 3.**
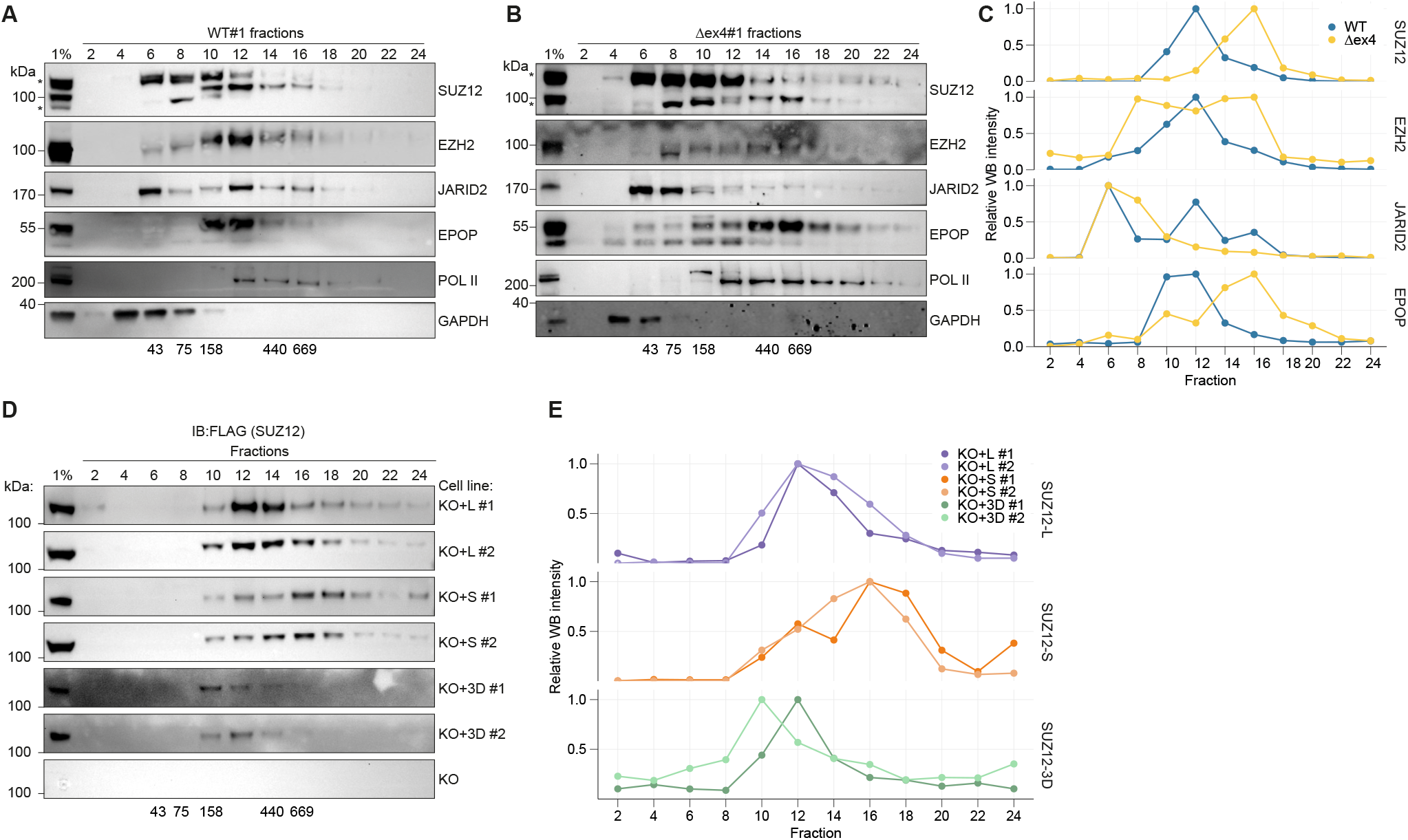
SUZ12-S favours PRC2 dimerization. (A,B) WB of glycerol gradient fractionation of WT (A) and Δex4 (B) nuclear extracts. (C) Quantification of WB bands in (A) and (B), normalised to highest intensity fraction. (D), WB of glycerol gradient fractionation of KO+L, KO+S and KO+3D cells. (E) Quantification of WB bands in (D) normalised to highest intensity fraction. In (A), (B), and (D), higher fractions correspond to higher molecular weights, numbers below gels correspond to expected fraction molecular weight (kDa), and asterisks indicate unspecific bands.

### SUZ12-L maintains global levels of H3K27 methylation

Next, we set out to investigate the contribution of each SUZ12 iso-form on PRC2 activity on chromatin using our different ESC lines. As an additional comparison for Δex4 cells, we included an ESC clone in which an unsuccessful CRISPR editing event resulted in a mutant allele with nearly constitutive splicing of exon 4 (herein, CSex4) and normal *Suz12* expression levels (Figures 4A and S4A). First, we profiled H3K27 bulk PTMs by WB and observed that, while CSex4 cells showed no major differences to control cells, Δex4 cells displayed lower levels of H3K27me2 and -me3, with higher levels of mono-methylation and acetylation (Figure 4A). Similar results were obtained when comparing SUZ12-L and SUZ12-S ESC rescue lines (Figure S2K). To validate these findings with an antibody-independent technique, and to additionally profile other histone PTMs, we analysed these cells with histone-MS. In line with the previous estimates ^25^, WT cells displayed approximately 85% of H3K27 methylation. This proportion was largely similar in CSex4 cells, while it dropped to ∼50% in Δex4 cells, with H3K27me2 and -me3 being the most affected modifications (Figures 4B and S4B). Notably, H3K27me2/3 loss in Δex4 cells occurred at histone peptides regardless of their H3K36 methylation status (Figure 4C). Moreover, the global H3K36 methylation rates were not affected in either CSex4 or Δex4 cells (Figure S4C), and no major changes in methylation or acetylation levels were observed at other residues (Table S3). Importantly, reintroduction of SUZ12-L in KO cells rescued a higher degree of methylation compared to SUZ12-S (Figures 4B and S4B). Overall, these results indicate that SUZ12-S alone is unable to maintain global physiological levels of H3K27 methylation; rather SUZ12-L is also necessary for this task.

**Figure 4.**
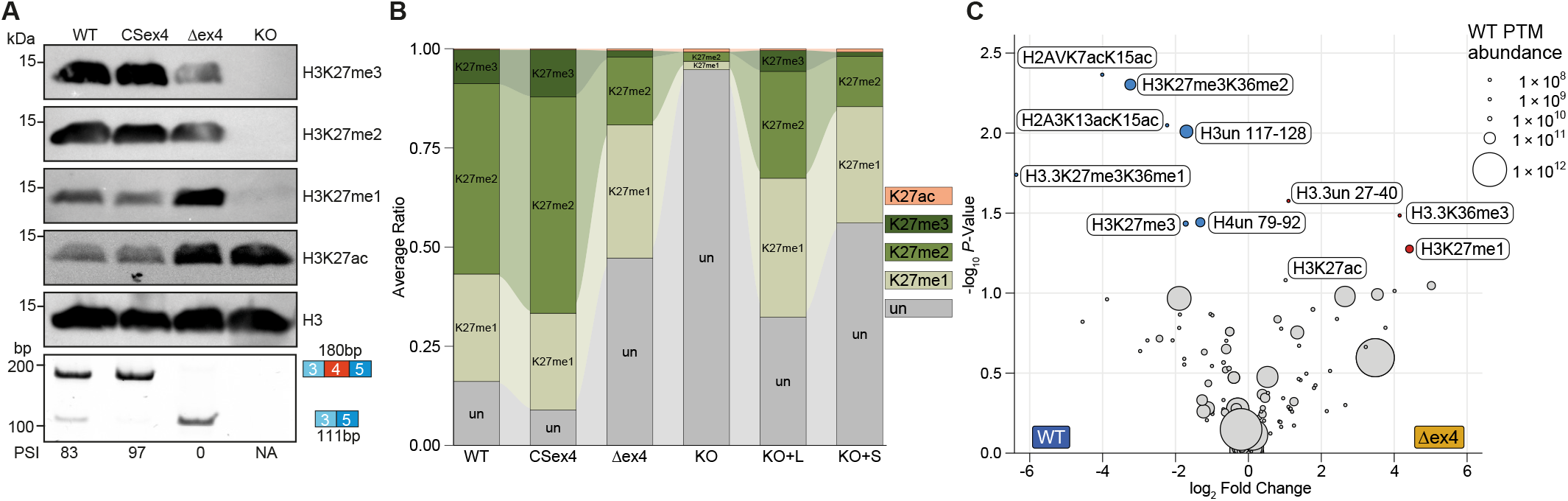
SUZ12-L is required for global H3K27 methylation. (A) Histone WB of H3K27 PTMs in ESCs (top); RT-RNA PCR showing the exon 4 splicing pattern in ESCs (bottom). The PSI is shown below the gel, and expected amplicon size, on the right. (B) Quantitative MS of histone H3K27 modifications in ESCs. The stacked barplot of the average PTM ratio of H3 (27-40) are shown. Analysis EpiProfile; combinatorial PTMs are deconvoluted and piled-up (see Methods). n = 3 clonal cell lines, un = unmodified. (C) Volcano plot of differentially abundant histone PTMs in WT and Δex4 ESCs. Significantly (adj. *P*-value *≤* 0.05) upregulated PTMs are shown in red, and significantly downregulated, in blue. Dot size corresponds to PTM abundance in WT samples. un = unmodified.

### SUZ12-S promotes highly localised H3K27me3 deposition

To assess whether the incorporation of SUZ12-L and SUZ12-S in PRC2 influences the genomic distribution of the complex, we analysed PRC2 chromatin occupancy and H3K27me3 deposition via ChIP-seq in WT, CSex4, Δex4 cells and KO cells (as a negative control). SUZ12 was found to occupy essentially the same set of promoters in WT, CSex4, Δex4 cell lines, suggesting that AS of exon 4 did not influence PRC2 target selection (Figures S5A and S5B). This is in line with previous reports showing that AEBP2 and JARID2 are dispensable for PRC2 targeting ^10,26^. However, CSex4 cells displayed reduced levels of H3K27me3 at target promoters. In contrast, these sites accumulated high levels of the mark in Δex4 cells (Figures S5B–S5D). Analysis of the distribution of H3K27me3 peaks in Δex4 cells revealed a lower proportion of intergenic peaks in favour of more promoter-proximal ones (Figure S5E), suggesting a redistribution of the mark towards canonical PRC2 targeting sites. To gain quantitative insights, we performed ChIP-seq using exogenous chromatin spike-in normalisation and obtained similar profiles (Figure 5A). We quantified global levels of H3K27me3 by assessing normalised read density over genomic bins of 2.5 kb in size (see Methods). This analysis confirmed that CSex4 cells display a similar distribution of H3K27me3 signal genome-wide with respect to WT cells, although with slightly lower intensity (Figure 5B). While most of the bins showed a reduction in H3K27me3 signal in Δex4 cells, the mark was selectively retained or even increased at bins encompassing a CGI targeted by SUZ12 (Figure 5C). Importantly, the drop in signal was already visible at bins flanking the targeted ones, suggesting that H3K27me3 deposition in these cells is more focalised (Figures 5C and 5D). In particular, analysis of the signal over CGIs revealed that Δex4 cells displayed higher and sharper peaks of H3K27me3 (Figure 5E). Altogether, these data indicate that SUZ12-S assembles a PRC2 complex capable of targeting and hyper-modifying PRC2 nucleation sites, but lacks the ability to spread the mark over extended genomic regions, for which SUZ12-L is necessary.

**Figure 5.**
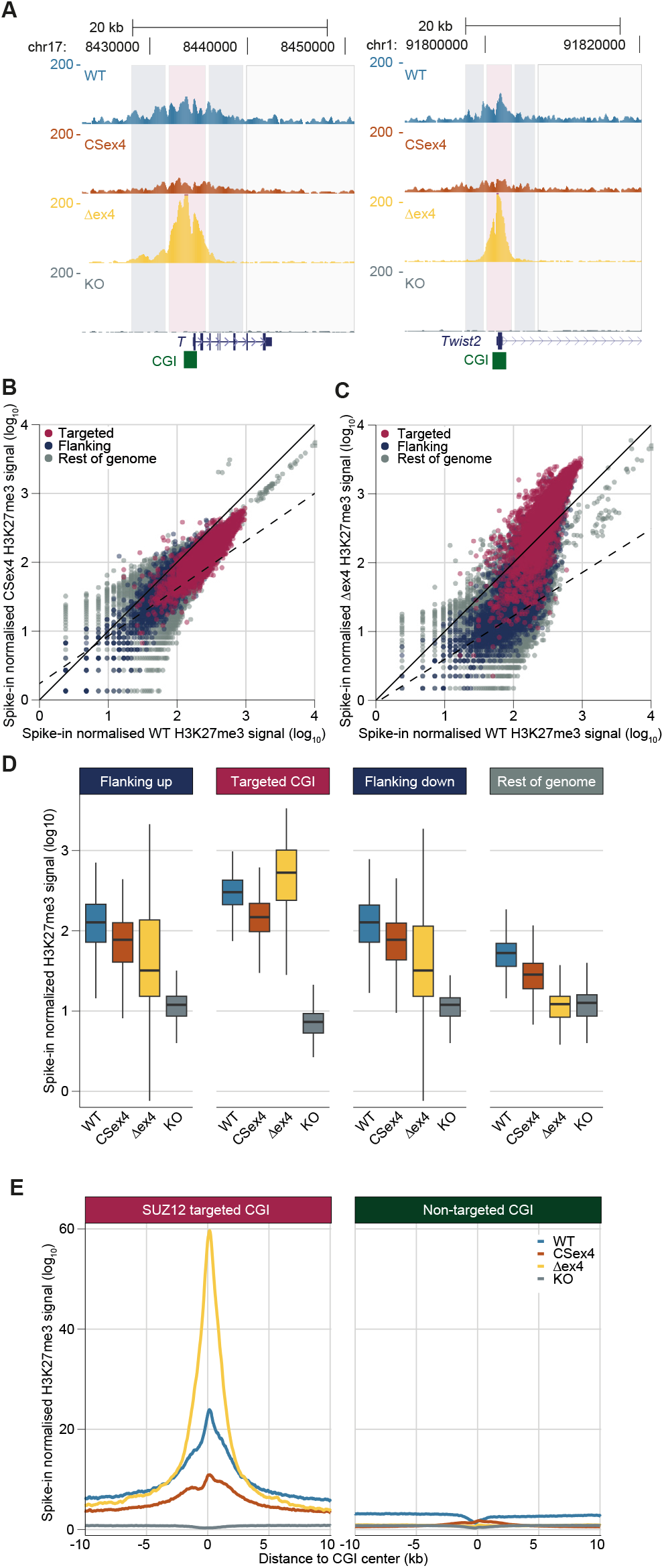
SUZ12-S deposits highly focalized H3K27me3 at PRC2 target promoters. (A) ChIP-seq tracks of H3K27me3 in two representative developmental genes targeted by PRC2. Locations of the CpG islands (CGIs) are indicated in green. (B,C) Scatter plot of spike-in normalised H3K27me3 signal in 2.5 kb genomic bins of WT vs CSex4 (B) or Δex4 (C). Bins overlapping SUZ12-targeted CGIs (n = 4189) are shown in amaranth; respective flanking bins (n = 5584), in dark blue; bins in the rest of the genome (n = 984638), in grey. Data from one clonal cell line. (D) Boxplots of spike-in normalised H3K27me3 signal in 2.5 kb genomic bins. Flanking up/down corresponds to the immediately upstream or downstream bin relative to the targeted CGI containing bin. The upper, centre and lower lines of the boxplot indicate 75%, 50% and 25% quantiles, respectively. Whiskers extend to the most extreme data point within 1.5× the interquartile range. (E), Metaplot of spike-in normalised H3K27me3 at SUZ12-targeted CGIs (left) or at other CGIs (right). Metaplots are centred on the middle of the CGI.

### SUZ12-S is necessary for maintaining PRC2-mediated repression at target sites

We next assessed how redistribution of H3K27 methylation affects gene expression. Transcriptomic analysis revealed that, despite the global reduction in H3K27me3 levels, Δex4 cells displayed no statistically differentially expressed genes (DEGs) (Figure 6A), suggesting that localised H3K27 hypermethylation of promoters is sufficient to maintain gene silencing. Surprisingly, CSex4 cells displayed 333 DEG (adj. *P*-value < 0.05), 71% of which were upregulated (Figure 6B). These genes were mostly associated with GO terms related to morphogenesis and tissue development (Figure S6A), in line with the role of PRC2 in silencing developmental genes in ESCs. As expected, full *Suz12* KO resulted in major transcriptomic dysregulation (Figure S6B). This was partially rescued by the reintroduction of either of the two isoforms (Figures S6C–S6E). However, the subset of rescued genes differed between the two isoforms (Figure 6C). Remarkably, PCR2 targets were only collectively and significantly upregulated in the absence of SUZ12-S, namely, in CSex4, KO+L and KO cells (Figures 6D and S6F).

**Figure 6.**
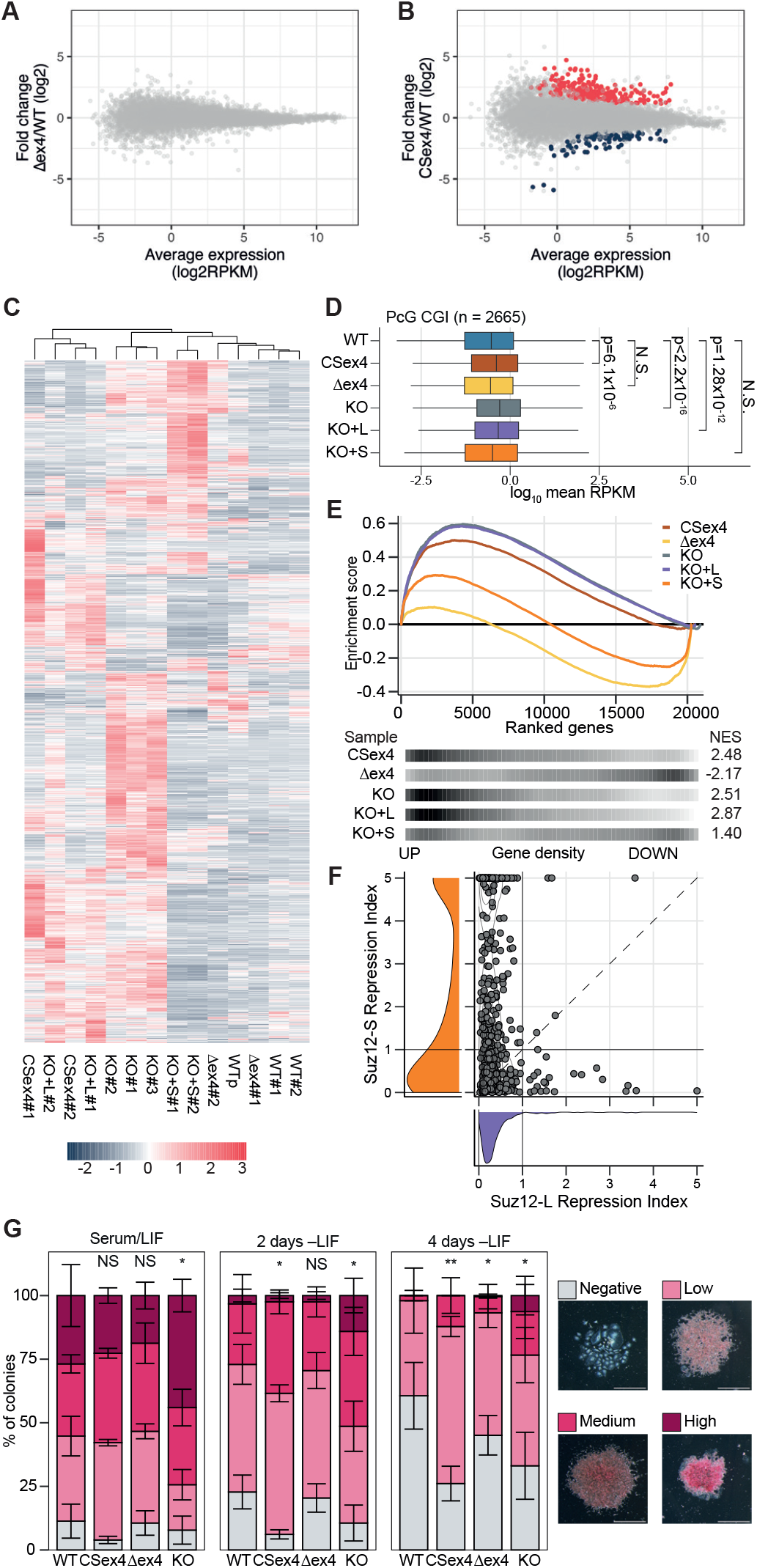
SUZ12-S but not SUZ12-L is sufficient to repress Polycomb target genes in ESCs. (A,B) Gene expression MA plots (log ratio vs abundance) of Δex4 (A) and CSex4 (B). Significantly upregulated DEGs are shown in red, and downregulated DEGs, in blue. Statistics, DESeq2. (C) Heatmap of 2052 SUZ12 CGI target genes across individual replicates coloured by RPKM z-score. (D) Boxplot of mean expression in ESCs of PRC2 target genes (n = 2665). Statistics, Wilcoxon test; *P*-value adjustment, Bonferroni; ns, not significant. (E) Plot showing positive or negative enrichment between gene ranks (DESeq2 stat) on the X-axis and SUZ12 CGI target genes, for different cell lines (top). Density distribution is shown for the ranked genes in the SUZ12 CGI target (bottom). Statistics, fgsea, FDR method, BH. (F) Scatterplot of SUZ12 gene expression repression index (see Methods) for SUZ12-L on the X-axis, and SUZ12-S, on the Y-axis. Each dot is a statistically upregulated DEG in the *Suz12* KO cells. Values close to zero correspond to expression levels comparable to those in *Suz12* KO cells; values close to 1, comparable to WT; and values above 1, more repressed than WT. Points above 5 are squished at 5. Dashed line, diagonal. Density distribution is shown on the left and bottom. (G) Stacked barplot quantification of alkaline phosphatase (AP) staining across three different timepoints of ESCs exiting pluripotency. Technical replicates (wells), n = 4; biological replicates (clonal cell lines), n = 2 or 3; error bars, sd. Statistics, Fisher’s exact test (see Methods); ns, not significant; *, significant in 50% of the tests; **, significant in more than 80% of the tests. Representative colonies are shown for each score (right); scale bar, 0.5 mm.

Consistently, gene set enrichment analysis (GSEA) revealed a significant upregulation of PRC2 target genes in CSex4 and KO+L cells, similar to KO cells, while an opposite trend was observed in Δex4 cells (Figure 6E). To further quantify these differences, we defined a *repression index* for each isoform, based on the strength of silencing of SUZ12 KO–upregulated genes, by each rescue (see Methods). Comparison of this index further supported that SUZ12-S was significantly (paired Wilcoxon test *P*-value < 2.2 ×10^*−*16^) more effective at silencing SUZ12 targets (Figure 6F). Taken together, these data demonstrate that SUZ12-S is necessary for proper Polycomb-mediated gene repression in ESCs. Lastly, we tested the impact of *Suz12* isoforms on ESC biology. In line with previous observations of *Suz12* KO cells ^27^, stemness, proliferation rates, and cell cycle profiles were not affected in either CSex4 or Δex4 cells as compared to control lines (Figure S6G–S6J). However, when challenged to differentiate through withdrawal of the leukaemia inhibitory factor (LIF), both CSex4 and KO cells (and to a lesser extent, Δex4 cells) displayed a delay in the exit from pluripotency (Figure 6G), a phenotype that has been associated with lack of Polycomb silencing in ESCs ^28,29^. This suggests that both *Suz12* isoforms are necessary for proper pluripotency exit.

## Discussion

We identified a novel placental mammals-specific *Suz12* isoform, *Suz12-S*, that is produced by exon 4 skipping (Figure 1). SUZ12-S protein favours PRC2.1 subtype assembly and increases its propensity to dimerise, resulting in higher levels of H3K27me3 deposition at target CGIs and stronger gene repression efficiency as compared to the longer, major isoform SUZ12-L.

Specific mutations of the SUZ12 ZnB and C2 domains have been shown to force the formation of either of the two PRC2 subtypes ^12^ and to determine their dimerisation propensities ^14^. Here, we propose that AS of exon 4, by modifying the length of the hinge connecting these two domains, serves as a physiological mechanism regulating both processes. By shortening the distance between the ZnB and C2 domains, exon 4 skipping disrupts the conformation required for binding to AEBP2 and JARID2^18,22^, hence favouring PRC2.1 formation (Figure 2). At the same time, closer proximity of the two domains prevents the C2-WD interaction on the same protomer, promoting domain swap–driven dimerisation ^14,15^ (Figure 3). However, as AEBP2 prevents dimerisation, it is also possible that loss of this protein from the SUZ12-S-containing PRC2 contributes to the increased propensity for dimerisation.

Previous work has shown a link between PRC2 dimerisation and H3K27 methylation at target promoters ^14^. We have now determined that SUZ12-S-containing PRC2 activity is strictly localised to CGIs and their nearest surroundings, suggesting higher affinity for these regions than SUZ12-L-containing PRC2 (Figure 5). Notably, the pattern of H3K27 methylation deposited by SUZ12-S is reminiscent of that of the ‘spreading-defective’ EED R302S mutant, in which H3K27me3 peaks corresponding to CGIs are maintained, while global levels of the mark drop significantly ^30,31^. Therefore, we propose that dimerisation may provide an opposite function with respect to the ‘read & write’ mechanism mediated by the EED–EZH2 axis, forcing local histone methyl-transferase activity and allowing for H3K27me3 nucleation sites to be established ^25^. Focalisation of PRC2 activity by SUZ12-S is likely independent of EZHIP-mediated PRC2 inhibition ^30,32^, as we did not detect significant interactions of SUZ12-S with this protein (Table S3).

Importantly, both *Suz12* isoforms are readily detected at the protein level and are co-expressed in virtually all cell and tissue types. Hence, our observations suggest that the WT landscape of H3K27 methylation results from the combined action of the two PRC2 isoforms with complementary functions: one by SUZ12-L, which accounts for the vast majority of H3K27 methylation genome wide, and the other one by SUZ12-S, exerting histone methyl-transferase activity in a highly localised manner at target promoters (Figures 4 and 5). Remarkably, this latter isoform is necessary and sufficient to maintain complete silencing of Polycomb target genes in mouse ESCs (Figure 6). The appearance of this isoform in placental mammals has therefore increased the repression toolkit in this clade, providing additional means to fine-tune gene regulation in the context of an ever-growing complexity of the regulatory landscape ^33^. Thus, exon 4 skipping exemplifies how PanAS events can contribute to gene functional diversification, particularly for TFs and chromatin regulators, in which they are strongly enriched ^17^. Specifically, in contrast to the more thoroughly studied AS ‘switches’, which generate distinct isoforms in different contexts ^34–36^, our data suggest that PanAS events can work as ‘splitters’. This allows SUZ12 to decouple global from local PRC2 activity across cell and tissue types. Therefore, through the interplay between two co-expressed, differentially specialised PanAS isoforms, genes can expand and diversify their functionality.

## Methods

### Alternative splicing analysis

RNA-seq AS analyses were performed using vast-tools v2.5.1^17^. Briefly, the software quantifies inclusion/exclusion levels of AS events, calculating the percentage of sequence inclusion (PSI) as: inclusion/(inclusion+exclusion) x 100. Quantification of the samples for human (*Homo sapiens*), mouse (*Mus musculus*), cow (*Bos taurus*), chicken (*Gallus gallus*), and zebrafish (*Danio rerio*) were retrieved from vastDB (vastdb.crg.eu). The samples of opossum (*Monodelphis domestica*) and elephant shark (*Callorhinchus milli*) were analysed previously ^37^. The PSI of SUZ12 exon 4 for manatee (*Trichechus manatus latirostri*s), elephant (*Loxodonta africana*), armadillo (*Dasypus novemcinctus*) and platypus (*Ornithorhynchus anatinus*) were quantified as follow: a 42-nt exon-exon junction ebwt-library for SUZ12 exons 3, 4, and 5 was generated using bowtie-build v1.1.2. The fastq files were trimmed to 50-nt sequences using vast-tools `Trim.pl` script, then aligned to the library using bowtie v1.1.2 with `-f -m 1 -v 2 ebwt-library`. The PSI was calculated from the alignment file, using the formula: sum of inclusion reads / (sum of inclusion reads + 2 x exclusion reads). The PSI of SUZ12 exon 4 in single cell human and mouse data and ENCODE mouse development were analysed with vast-tools. All analysed samples were publicly downloaded from SRA and are listed in Table S1. For PSI validation, RNA extraction was performed using the RNeasy kit (Qiagen, 74134) according to the manufacturer’s instructions. Retro-transcription (RT) was performed using SuperScript III Reverse Transcriptase (Life Technologies, 18080-085) using anchored oligo-dT. cDNA was PCR amplified using primers annealing in exon 3 and exon 5 (Table S2). Amplicons were resolved on 7.6% TBE-acrylamide gels (PanReac Applichem, A1672) at 210V for 20 min and stained with 1:10000 SBYR Safe in H_2_O (Life Technologies, S33102) for 3 min. Images were acquired with the GelDoc XR+ system (Bio-Rad) using non saturating settings. Band intensity was quantified using Image Lab v6.0 (Bio-Rad) with background subtraction of 18mm and PSI was quantified as: inclusion / (inclusion + exclusion) ×100.

### Phylogenetic sequence comparison

SUZ12 exon 4 conservation and sequence logo analysis is described in detail at github.com/Ni-Ar/msbt. Briefly, SUZ12 exon 4 coordinates are manually curated in 153 species genome assemblies. The DNA sequences were retrieved with the `coord2seq.sh` script. The output was plotted with custom-made scripts in R using ggseqlogo ^38^.

### Structural analysis

PRC2 structures (PDB: 6WKR ^7^; 6NQ3^14^) were analysed and visualised with ChimeraX v1.5^39^.

### Cell lines

Mouse embryonic stem cells (ESCs) from a 129xC57BL/6J background ^40^ were routinely cultured on 0.1% gelatin-coated tissue culture plates in Glasgow Modified Eagle’s Medium (Sigma, G5154), supplemented with 10% inactivated foetal bovine serum (Cytiva HyClone SV30160.03), Glutamax (Gibco), penicillin/streptomycin (Gibco), non-essential amino acids (Gibco), β-mercaptoethanol (Gibco) and leukaemia inhibiting factor (LIF; produced in-house). Derived cell lines were generated as described in Figure S2a,h. Briefly, Δex4 cells were generated by CRISPR-Cas9-mediated deletion of the genomic region encompassing exon 4 and part of the flanking intronic sequences. Similarly, *Suz12* KO cells were generated by deleting exons 2 and 3, hence generating a frameshift in the remaining transcript (regardless of the inclusion or skipping of exon 4). For the isoform-specific rescues of *Suz12*, a cassette including the CAG promoter driving the expression of *Suz12-L* or *Suz12-S* cDNA fused to a T2A:EGFP for FACS-mediated clones selection was inserted via CRISPR-Cas9 knock-in inside the Rosa26 locus of KO cells. Genomic editing of each clonal line was confirmed via Sanger sequencing. All guide RNAs and genotyping primers used are listed in Table S2.

### RT-qPCR

RNA extraction and RT were performed as described above. qPCR was performed in Lightcycler 480 (Roche, 05015243001) using the NZYSpeedy qPCR Green Master Mix (2x) (NZYtech, MB224) using 4ng of cDNA per well. Primer sequences are reported in Table S2.

### Protein immunoprecipitation

To isolate the nuclei, cells were resuspended in a hypotonic buffer (10 mM Tris-HCl pH 7.4, 10 mM KCl, 15 mM MgCl_2_) and incubated for 10 min on a rotating wheel at 4°C. Nuclei were then pelleted by centrifugation at 700 g for 5 min at 4°C. Nuclei pellet was then resuspended in nuclear lysis buffer (150 mM NaCl, 50 mM Hepes pH 7.5, 2.5 mM MgCl_2_, 0.5% NP-40) supplemented with 100 U/ml Benzonase (Novagen, 71205) and protease and phosphatase inhibitors, and incubated for 1 h at 4°C on a rotating wheel. The extract was then clarified by centrifugation at max speed for 10 min in a tabletop centrifuge at 4°C. Protein concentrations were quantified with the BCA assay. Per each IP, 1 to 5 mg of protein were diluted at least 1:3 in the IP buffer (150 mM NaCl, 50 mM Hepes pH 7.5, 0.5% NP-40, 5 mM EDTA) supplemented with protease/phosphatase inhibitors. To this, 2 μg/ml of antibody and 20 μL/mL of Protein A Dynabeads (Invitrogen, 10002D) were added and incubated overnight at 4°C on a rotating wheel. Beads were washed with the IP buffer three times. For WB, proteins were eluted in the loading buffer for 15 min at 70°C and loaded on SDS-PAGE. For MS application, on-beads digestion was performed as follows: beads were washed three times using ABC buffer (200 mM ammonium bicarbonate) and resuspended in 6 M urea in ABC. Samples were then reduced with DTT (30 nmol, 37°C, 60 min), alkylated in the dark with iodoacetamide (60 nmol, 25°C, 30 min), diluted with ABC to a final concentration of 1 M urea and digested using trypsin (Promega, V5113; 3 μg/mL, 37°C, 16 h). After digestion, the peptide mix was acidified using formic acid and desalted with a MicroSpin C18 column (The Nest Group) prior to MS analysis.

### Targeted proteomics

Samples were analysed using an Orbitrap Eclipse (Thermo Fisher Scientific) coupled to an EASY-nanoLC 1200 UPLC system (Thermo Fisher Scientific). Peptides were loaded directly onto the analytical column and were separated by reversed-phase chromatography using a 50-cm column with an inner diameter of 75 μm, packed with 2 μm C18 particles spectrometer (Thermo Scientific). Chromatographic gradients started at 95% buffer A (0.1% formic acid) and 5% buffer B (0.1% formic acid, 80% acetonitrile) with a flow rate of 300 nL/min for 5 min and gradually increased to 25% buffer B and 75% A in 52 min and then to 40% buffer B and 60% A in 8 min. After each analysis, the column was washed for 10 min with 10% buffer A and 90% buffer B. The mass spectrometer was operated in positive ionisation mode with an EASY-Spray nanosource at 2.4kV and at a source temperature of 305°C. Full MS scans with 1 micro scan at resolution of 30,000 were used over a mass range of m/z 350-1400 with detection in the Orbitrap mass analyzer. A parallel reaction monitoring (PRM) method was used for data acquisition with a quadrupole isolation window set to 1.4 m/z and MS-MS scans over a mass range of m/z 300-2000, with detection in the Orbitrap at resolution of 60,000. MS-MS fragmentation was performed using HCD at 30 NCE, the auto gain control (AGC) was set to 1 ×10^5^ and maximum injection time of 118 ms. Peptide masses (m/z) were defined in the mass list table (Table S3) for further fragmentation. Skyline v20.2^41^ was used to extract the fragment areas of each peptide. The sum of the fragments area of SUZ12-L specific peptide (TFKVDDMLSKVEK) and SUZ12-S specific (SLSAHLQLTFTGFFHK) were normalised to a common peptide (`NLIAPIFLHR`) area and plotted in R.

### IP-MS

Samples were analysed using a LTQ-Orbitrap Velos Pro mass spectrometer (Thermo Fisher Scientific) coupled to an EASY-nLC 1000 (Thermo Fisher Scientific (Proxeon), Odense, Denmark). Peptides were loaded onto the 2-cm Nano Trap column with an inner diameter of 100 μm packed with C18 particles of 5 μm particle size (Thermo Fisher Scientific) and were separated by reversed-phase chromatography using a 25-cm column with an inner diameter of 75 μm, packed with 1.9 μm C18 particles (Nikkyo Technos Co., Ltd.). Chromatographic gradients started at 93% buffer A and 7% buffer B with a flow rate of 250 nl/min for 5 min and gradually increased 65% buffer A and 35% buffer B in 60 min. After each analysis, the column was washed for 15 min with 10% buffer A and 90% buffer B. The mass spectrometer was operated in positive ionisation mode with nanospray voltage set at 2.1 kV and source temperature at 300°C. Ultramark 1621 was used for the external calibration of the FT mass analyser prior the analyses and an internal calibration was performed using the background polysiloxane ion signal at m/z 445.1200. The acquisition was performed in data-dependent acquisition (DDA) mode and full

MS scans with 1 micro scan at resolution of 60,000 were used over a mass range of m/z 350-2000 with detection in the Orbitrap. AGC was set to 1×10^6^, dynamic exclusion (60 s) and charge state filtering disqualifying singly charged peptides was activated. In each cycle of DDA analysis, following each survey scan, the top twenty most intense ions with multiple charged ions above a threshold ion count of 5000 were selected for fragmentation. Fragment ion spectra were produced via collision-induced dissociation (CID) at normalised collision energy of 35% and they were acquired in the ion trap mass analyzer. AGC was set to 1000, and an isolation window of 2.0 m/z, an activation time of 10 ms and a maximum injection time of 100 ms were used. All data were acquired with Xcalibur software v2.2. Digested bovine serum albumin (New England Biolabs, P8108S) was analysed between each sample to avoid sample carryover and to assure stability of the instrument and QCloud ^42^ has been used to control instrument longitudinal performance during the project. Acquired spectra were analysed using the nextflow ^43^ pipeline `proteomicslfq` v1.0.0 (nf-co.re/proteomicslfq). Briefly, Thermo raw files were searched against the mouse UniProt protein database version `2022_03` containing both canonical and alternative isoforms and common contaminants with MSGF+ and Comet. Results between the two search engines were combined with OpenMS’ ConsensusID. Label free protein intensities were quantified using spectral counting of Peptide Spectral Matches (PSMs) with options:`-quantification_method ‘spectral_counting’`; `-protein_quant ‘shared_peptides’` (quantify proteins based on peptides mapping to single and multiple proteins too, but only mapping peptides greedily for its best group by inference score); `-protein_inference ‘bayesian’` (inference to group proteins is controlled by calculating scores at the protein group level and to potentially modify associations from peptides to proteins using Bayesian statistics). The resulting output mzTab file was then processed using the DEP R package ^44^ to perform differential enrichment analysis by applying empirical Bayes statistics on protein-wise linear models. Prey-to-bait stoichiometric ratio was defined as follows: PSMs for each protein were normalised to its length and divided by the corresponding average value of SUZ12 in each genotype.

### Chromatin fractionation

Chromatin fractionation protocol was adapted from ^45^. Briefly, nuclei were isolated by resuspending the cells in Buffer A (10 mM Hepes pH 7.9, 10 mM KCl, 1.5 mM MgCl_2_, 0.34 M Sucrose, 10% Glycerol, 1 mM DTT, and 0.1% Triton) supplemented with protease and phosphatase inhibitors. Nuclei were separated from the supernatant (1300 g, 4°C, 5 min) and washed again in Buffer A. After that, nuclei were resuspended in Buffer B (3 mM EDTA, 0.2 mM EGTA, 1 mM DTT) supplemented with protease and phosphatase inhibitors, to induce breakdown of the nuclear membrane. The insoluble chromatin fraction was spun down (1700 g, 4°C, 5 min), washed once in Buffer B and then resuspended in SDS buffer (50 mM Tris-HCl pH 7.4, 4 mM EDTA pH 8.0, 2% SDS). Samples were boiled at 98°C for 10 min and chromatin was sonicated to allow complete resuspension.

### Protein complex fractionation

Nuclear extracts were obtained as described for the protein IP protocol and then concentrated to a final volume of 500 μL by centrifugation in a 5 kDa Amicon Ultra filter unit (Millipore, UFC900508) at 4000 rpm at 4°C. The protein weight ladder was obtained using the Gel Filtration HMW Calibration Kit (Cytiva, 28403842) according to manufacturer’s instructions to obtain a solution of 500 μL containing 0.8 mg Ovalbumin (43 kDa, monomer), 0.5 mg Conalbumin (75 kDa, monomer), 0.8 mg Aldolase (158 kDa, tetramer), 4.5 ng Ferritin (440 kDa, 24-mer), and 1.25 mg Thyroglobulin (669 kDa, dimer). Prior to loading, both the nuclear extract and the protein ladder were diluted 1:4 in the dilution buffer (150 mM KCl, 0.2 mM EDTA, 0.1% NP-40, 20 mM Hepes pH 7.9, 0.5 mM DTT). The glycerol gradient was prepared as follows: 5 ml of a 10% Glycerol solution (150 mM KCl, 0.2 mM EDTA pH 8.0, 0.1% NP-40, 20 mM Hepes pH 7.9, 10% Glycerol) were overlaid on the same volume of a 50% Glycerol solution (150 mM KCl, 0.2 mM EDTA pH 8.0, 0.1% NP-40, 20 mM Hepes pH 7.9, 50% Glycerol) in an Ultra Clear 13.2 mL tube (Beckman Coulter, 344059), and mixed using a Gradient Master 107 IP (Biocomp). Gradients were incubated at 4°C for 2 h prior to centrifugation. To separate the proteins, the extract and the protein ladder were overlaid onto separate gradients and centrifuged with an SW41Ti rotor (Beckman Coulter, 20U113807) in an Optima XPN-100 Ultracentrifuge (Beckman Coulter) at 33000 rpm at 4°C for 22 h. After centrifugation, 24 fractions of 500 μL were collected, starting from the top of the tube. For Western blot analysis, 5% of each fraction was used. As a weight reference, fractions of the protein ladder gradient were run on a NuPAGE gel, stained using BlueSafe (Nzytech, MB15201) (Figure S7A-S7B).

### Histone MS

Histone proteins were extracted as described in ^46^. For each cell line (n = 3 clonal cell lines) 20 million ESCs were harvested and washed twice in PBS and stored at -70°C until processing. Frozen cell pellets were resuspended in 300 μL of cold hypotonic buffer (10 mM Tris-HCl pH 8, 1.5 mM MgCl_2_, 1 mM KCl, 10X PhosphoSTOP (Roche, 4906845001), 2X Protease Inhibitor Cocktail (Roche, 11836170001), 1 mM DTT, 0.5 mM PMSF). Samples were lysed on a rotating wheel for 30 min and spun down at 10000 g for 10 min at 4°C. Cell pellets were resuspended in 1300 μL of 0.2 M H_2_SO_4_ and incubated overnight at 4°C. Samples were spun down at 16000 g for 10 min at 4°C, supernatant was transferred to a new tube and TCA was added dropwise to a final concentration of 33% to precipitate soluble histones. After 2 h incubation on ice samples were spun down at 16000 g for 10 min. Histone pellets were washed twice with 1 mL of ice cold acetone and air dried for 2 min before resuspending in 100 μL H_2_O. Purified histone proteins were quantified with BCA, diluted to 1 μg/μL and 10μg of each sample were chemically derivatized with propionic anhydride, digested with trypsin and derivatized again with propionic anhydride. Briefly, samples were dissolved in 9 μL of H_2_O and 1 μL of triethyl ammonium bicarbonate was added to bring the pH to 8.5. Propionylation was carried out by adding 1 μL of a propionic acid solution (1% propionic anhydride, Sigma, 240311) with vortexing and incubation for 2 min. The reaction was quenched with 1 μL of 80 mM hydroxylamine (Sigma, 467804) and samples were incubated at room temperature for 20 min. Tryptic digestion was performed for 3 h with 0.1 μg trypsin at 37°C (Promega, V5113) per sample. Digested peptides were propionylated again with 3 μL of propionic acid solution with vortexing and incubation for 2 min. The reaction was quenched with 1 μL of 80 mM hydroxylamine and samples were incubated at room temperature for 20 min. Samples were acidified by adding 50 μL of 5% formic acid, vacuum dried and desalted with C18 ultramicrospin columns (The Nest Group). A 2 μg aliquot of the peptide mixture were analysed using a LTQ-Orbitrap Fusion Lumos mass spectrometer (Thermo Fisher Scientific) coupled to an EASY-nLC 1200 (Thermo Fisher Scientific) in both data independent acquisition (DIA) and data dependent acquisition (DDA) methods. Peptides were loaded directly onto the analytical column and were separated by reversed-phase chromatography using a 50-cm column with an inner diameter of 75 μm, packed with 2 μm C18 particles spectrometer (Thermo Scientific) with a 60 min chromatographic. The mass spectrometer was operated in positive ionisation mode on both DDA and DIA modes. The DDA method was driven by the “Top Speed” acquisition algorithm, which determined the number of selected precursor ions for fragmentation, whilst the DIA method consisted in repetitive acquisition cycles of 25 staggered windows of 24 m/z covering a mass range from 350 to 1850 m/z. DIA runs were analysed using EpiProfile v2.1^47^ with parameters: `norganism=2`, `nsource=1`, `nsubtype=0`, `def_ptol=10`, `soutput=‘11’`. The output was analysed using the R packages `niar` (github.com/Ni-Ar/niar) to process and visualise the PTM ratios and DEP ^44^ to perform PTM differential enrichment analysis using empirical Bayes statistics. In addition, to validate the results, DDA runs were analysed using the Proteome Discoverer software suite v2.5 (Thermo Fisher Scientific), and the Mascot ^48^ search engine v2.6 (Matrix Science) was used for peptide identification. Data were searched against the SwissProt mouse database plus the most common contaminants. A first search was done considering propionylation on N-terminal and lysines as variable modifications. With the proteins obtained in this search, a new database was generated, and a second database search was done considering propionylation on N-terminal as fixed modification and propionylation on lysines, dimethyl lysine, trimethyl lysine, propionyl + methyl lysine, and acetyl lysine as variable modifications. Precursor ion mass tolerance of 7 ppm at the MS1 level was used, and up to 5 missed cleavages for trypsin were allowed. False discovery rate in peptide identification was set to a maximum of 5%. The results from the Proteome Discoverer software were used to generate a library in Skyline ^41^ v22.2.1.306 that was used to detect, extract and quantify the MS1 and MS2 signal of the histone peptides and their variants from the DIA runs.

### Chromatin immunoprecipitation

Cells were cross-linked by incubation with 1% formaldehyde (Sigma, F8775) in PBS for 10 min at room temperature. Glycine was added to a final concentration of 125 mM and incubated for an additional 5 min to stop cross-linking. Cells were washed twice with PBS and then collected using a cell lifter (Corning, 3008) in PBS supplemented with protease inhibitors. Pellets were snapfrozen in dry ice and stored at -70°C until use. ChIP was performed as follows: lysis was induced by resuspending the cells in Lysis/IP buffer composed of 1 volume of SDS buffer (100 mM NaCl, 50 mM Tris-HCl pH 8.0, 5 mM EDTA pH 8.0, 0.5% SDS) and 0.5 volumes of Triton buffer (100 mM NaCl, 100 mM Tris-HCl pH 8.6, 5 mM EDTA pH 8.0, 5% Triton) supplemented with protease and phosphatase inhibitors. Chromatin was sonicated using a Bioruptor Pico (Diagenode) in 15 ml sonication tubes (Diagenode, C01020031) using 350 mg/mL of sonication beads (Diagenode, C03070001) for 40 cycles of 30 s ON, 30 s OFF. Per each ChIP sample, a chromatin amount equivalent to 30 μg of DNA was used. IP was performed by addition of 4 μg/mL of antibody, and 20 μL/mL of Protein A Dynabeads and incubation overnight at 4°C on a rotating wheel. When used, spike-in normalisation was carried out according to guidelines published in ^49^ Briefly, a 2.5% equivalent of sonicated drosophila chromatin (prepared in-house) was added, together with 1 μg/mL of Spike-In Antibody. The day after, the beads were washed three times with the Dilution buffer. Chromatin was then eluted by resuspending the beads in elution buffer (0.1 M NaHCO_3_, 1% SDS) and incubating at 65°C for 30 min in a Thermomixer shaking at 1000 rpm. Eluted chromatin was then de-crosslinked by addition of NaCl to a final concentration of 200 mM and incubation at 65°C overnight. Protein digestion was carried out by addition of Proteinase K to a final concentration of 0.1 mg/mL, EDTA to a final concentration of 10 mM, and Tris-HCl pH 6.5 to a final concentration of 40 mM and incubated at 45°C for 1 h. DNA was purified using the PCR Purification Kit (Qiagen, 28106) and eluted in ultrapure H_2_O. ChIP was performed in biological duplicates except for spike-in normalised H3K27me3 ChIP-seq, which was performed only in one clonal line per genotype. For ChIP-qPCR, 1% of the eluted DNA was used per well. qPCR was performed as described above. For ChIP-seq, DNA was quantified using Qubit (Thermo Fisher Scientific) and libraries were prepared using the NEBNext Ultra DNA Library Prep (Illumina, E7370) according to the manufacturer’s protocol. Briefly, starting amount of 4 ng of input and ChIP enriched DNA were subjected to end repair and addition of ‘A’ bases to 3’ ends, ligation of adapters and USER excision. Purification steps were performed using AgenCourt AMPure XP beads (Beckman Coulter, A63882). Library amplification was performed by PCR using NEBNext Multiplex Oligos for Illumina. Final libraries were analysed using Agilent Bioanalyzer to estimate the concentration and size and were then quantified by qPCR using the KAPA Library Quantification Kit prior to amplification with Illumina’s cBot. Libraries were sequenced on a single end for 50+8bp on Illumina’s HiSeq2500. A minimum of 30 ×10^6^ reads per sample was generated. ChIP-seq samples including spike-in were mapped against a synthetic genome constituted by the mouse and the drosophila chromosomes (mm10 + dm3) using Bowtie with the option `-m 1` to discard reads that did not map uniquely to one region ^50^. Peak calling was carried out by running MACS on each replicate individually, with the default parameters but with the shift-size adjusted to 100 bp to perform the peak calling against the corresponding control sample ^51^. DiffBind ^52^ was run next over the union of peaks from each pair of replicates of the same experiment to find those peaks that were significantly enriched in both replicates in comparison to the corresponding controls (DiffBind 2.0 arguments: `categories = DBA_CONDITION`, `block = DBA_REPLICATE` and `method` `= DBA_DESEQ2_BLOCK`). In all cases, DiffBind peaks with *P*-value < 0.05 and FDR < 1 ×10^*−*5^ were selected for further analysis. The genome distribution of each set of peaks was calculated with SeqCode ^49^ by counting the number of peaks fitted on each class of region according to RefSeq ^53^ annotations. Promoter is defined as the region between 2.5 kbp upstream and 2.5 kbp downstream of the transcription start site (TSS). Genic regions correspond to the rest of the gene (the part that is not classified as promoter) and the rest of the genome is considered as intergenic. Peaks that overlapped with more than one genomic feature were proportionally counted the same number of times. CpG islands (CGIs) were retrieved from the UCSC genome browser ^54^. To generate our collection of CGIs associated with PRC2, we performed the overlap of our consolidated set of SUZ12 peaks against the UCSC CpG track using SeqCode. The mm10 genome was segmented into bins of 2.5 kbp and those bins overlapping at least one nucleotide with a PRC2 CGI were considered to be bins targeted by SUZ12, while the rest constituted our set of non-targeted bins. Target genes for each sample were retrieved using SeqCode by matching the ChIP-seq peaks in the region 2.5 kbp upstream of the TSS until the end of the transcripts as annotated in RefSeq. Genome-wide profiles, meta-plots and values for boxplots/scatterplots of quantification of ChIP-seq signal strength were generated with SeqCode, instructing the software to normalise each value appropriately for the corresponding number of mouse or spike-in reads of the experiment. Screenshots of ChIP-seq tracks were generated using the UCSC genome browser. Tracks of replicates were grouped and merged using the ‘transparent’ overlay option, using a windowing function with the ‘maximum’ settings and a smoothing window of two pixels.

### RNA-seq analysis

RNA extraction was performed as described above. Sequencing libraries were prepared using the TruSeq stranded mRNA Library Prep (Illumina, 20020595) according to the manufacturer’s protocol. Briefly, 1000 ng of total RNA was used for poly(A)-mRNA selection using poly-T oligo attached magnetic beads using two rounds of purification. During the second elution of the poly-A RNA, the RNA was fragmented under elevated temperature and primed with random hexamers for cDNA synthesis. Then, the cleaved RNA fragments were copied into first strand cDNA using the SuperScript II reverse transcriptase (Invitrogen, 18064-014) and random primers. After that, a second strand cDNA was synthesised, removing the RNA template and synthesising a replacement strand, incorporating dUTP in place of dTTP to generate ds cDNA using DNA Polymerase I and RNase H. These cDNA fragments, then had the addition of a single ‘A’ base to the 3’ ends of the blunt fragments to prevent them from ligating to one another during the adapter ligation. A corresponding single T nucleotide on the 3’ end of the adapter provides a complementary overhang for ligating the adapter to the fragments. Subsequent ligation of the multiple indexing adapter to the ends of the ds cDNA was done. Finally, PCR selectively enriched those DNA fragments that had adapter molecules on both ends. The PCR was performed with a PCR Primer Cocktail that anneals to the ends of the adapters. Final libraries were analysed using Bioanalyzer DNA 1000 (Agilent, 5067-1504) to estimate the quantity and validate the size distribution, and were then quantified by qPCR using the KAPA Library Quantification Kit KK4835 (Roche, 07960204001) prior to the amplification with Illumina’s cBot. Libraries were sequenced on a single end for 50+8bp on Illumina’s HiSeq2500. A minimum of 40× 10^6^ reads per sample was generated. RNA-seq samples were mapped against the mm10 mouse genome assembly using TopHat ^55^ with the option `-g 1` to discard those reads that could not be uniquely mapped in just one region. DESeq2^56^ was run to quantify the expression of every annotated transcript using the Ref-Seq catalogue of exons ^53^ and to identify each set of differentially expressed genes between the two conditions. Analysis of GO-term enrichment was performed using the Web-based Gene Set Analysis Toolkit ^57^. The ‘Overlap’ measure represents the number of genes in common between the queried gene set and the one represented, while the ‘Enrichment ratio’ is the ratio between the observed and the expected overlap. This measure was used to rank entries in the plot. Only entries with an FDR < 0.01 were reported. Gene set enrichment analysis was performed in R using the fgsea package ^58^ with `eps = 0`, `nPermSimple = 1000`, `proc = 4`. Results were plotted with custom made script in R. SUZ12 *repression index* was calculated for all upregulated DEGs in *SUZ12* KO using log_2_ normalised DESeq2 counts averaged between replicates as follow: (Rescue - WT) / (KO - WT), where the numerator represents the difference in gene expression relative to WT for either KO+S or KO+L and the denominator normalise for how much a gene is up-regulated in the KO.

### Cell differentiation

Cells were plated in serum/LIF conditions to clonal density (10^2^/cm^2^). Differentiation was triggered by LIF withdrawal for 48 or 96 h. After a total of five days of growth, cell pluripotency was evaluated by assessing alkaline phosphatase (AP) activity using the leukocyte alkaline phosphatase kit (Sigma, 86R-1KT) according to manufacturer’s instructions. Colonies were scored as high, medium, low or negative according to the staining intensity. A minimum of 70 colonies were counted for each replicate (technical replicates, n = 4 wells; biological replicates, n = 2 clonal cell lines; timepoints, n = 3 days). Statistical significance was calculated using Fisher’s exact test.

### Self renewal and proliferation capacity assessment

Cells were pulse-labelled with 10 μM BrdU (BD Biosciences, 51-2420KC) for 30 min, then harvested and fixed in 75% ethanol at-20°Cfor at least 1 h. Cells were resuspended in 2 N HCl with 0.5% Triton X-100 to denature the DNA and then resuspended in 0.1 M Na_2_B_4_O_7_ 10 H_2_O, pH 8.5 to neutralise the acid. BrdU was stained using a GFP-conjugated anti-BrdU antibody (BD Biosciences, 347583) according to manufacturer’s instructions. Finally, DNA content was assessed via propidium iodide (PI) incorporation by resuspending the cells in a PBS solution containing 15 μg/mL PI, 1 mM sodium citrate, and 0.01 mg/mL Ribonuclease A over night, prior to flow cytometry analysis. Flow cytometry analysis was performed using the LSR II Flow Cytometer (BD, H48200055). Data was collected using the FACSDiva Software (BD) v6.1.3 (BD LSR II). Analysis was carried out using Flowjo (BD) v10.8.2. Gating strategy is described in Figure S7C. For doubling time assessment, cells were counted and re-plated every other day. For pluripotency assessment, cells were stained for SSEA1 (Invitrogen, 50-8813-42) and analysed as described above. Gating strategy is described in Figure S7D.

## Code

All the code used to generate the results is available at the GitHub repository github.com/Ni-Ar/SUZ12-AS-project (*work in progress*)

## Data

ChIP-seq and RNA-seq fastq files and processed data are deposited on GEO with accession GSE223666. Proteomic data is deposited on PRIDE with accession: PXD000000 (*work in progress*).

## ACKNOWLEDGEMENTS

We are thankful to Barbara Pernaute and Sergi Aranda for fruitful discussion and suggestions. RNA samples from zebrafish developmental stages were kindly provided by Jon Permanyer. Human organoids samples were generated by the Tissue Engineering Unit of the CRG. Proteomics samples were generated in the CRG/UPF Proteomics Unit which is part of the Proteored PRB3 and is supported by grant PT17/0019, of the PE I+D+i 2013-2016, funded by ISCIII and ERDF. We thank Zuo-Fei Yuan (University of Pennsylvania) and Simone Sidoli (Albert Einstein College of Medicine) for fruitful discussion on histone PTMs analysis and Zuo-Fei Yuan for giving us early access to EpiProfile v2.1. Niccolò Arecco was supported by the EMBO Postdoctoral Fellowship (ALTF 695-2019) and by the Spanish Ministry of Science and Innovation Juan de la Cierva Formación (FJC2018-038498-I). Ivano Mocavini was supported by an FPI fellowship from the Spanish Ministry of Science and Innovation.

## Supplementary Figures

**Supplementary Figure 1.**
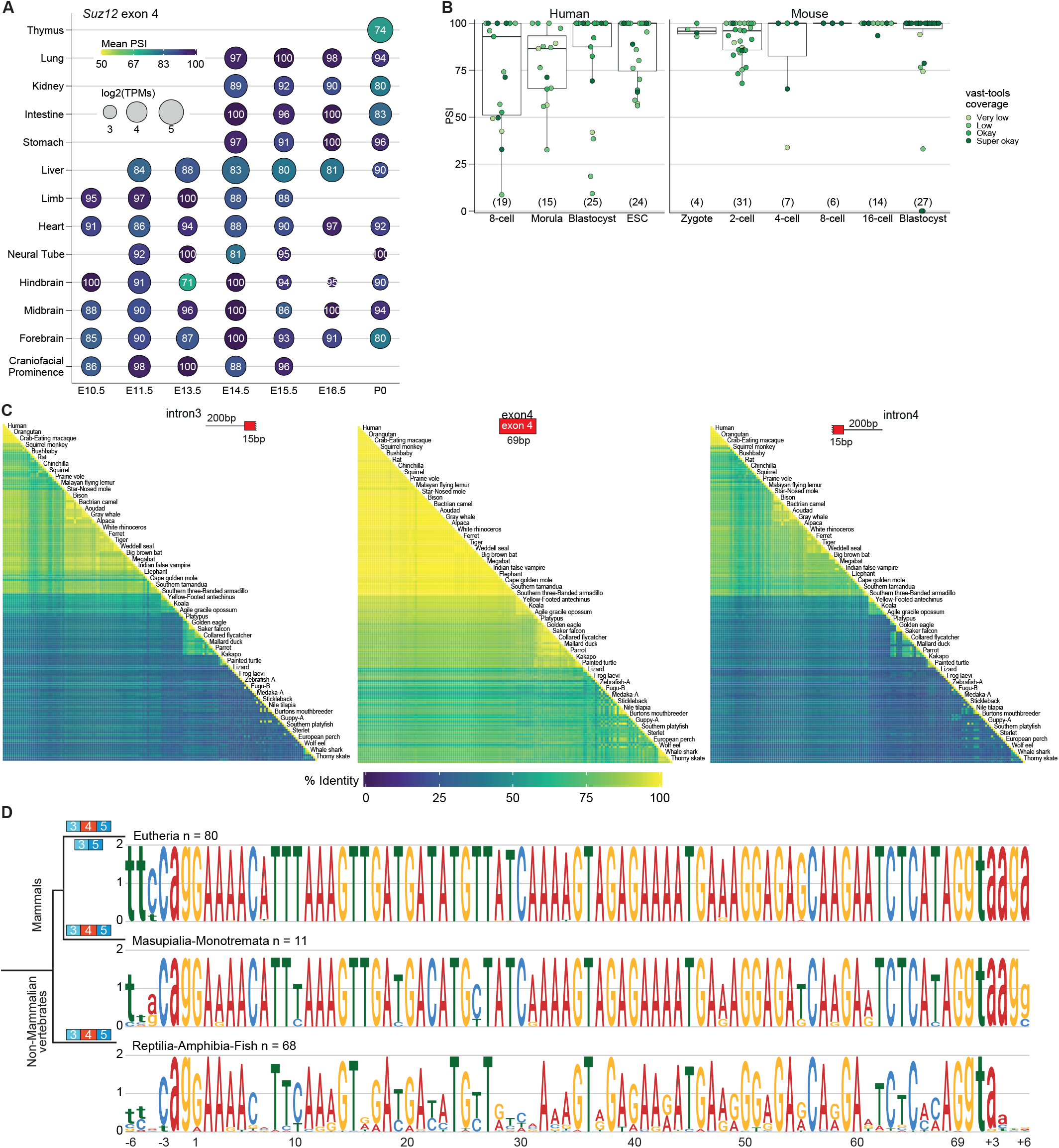
*Suz12* exon 4 sequence inclusion, and sequence conservation across vertebrates. (A), ENCODE mouse development samples quantification of Suz12 gene expression (dot sizes) and exon 4 PSI (dot colours). Numbers inside the dots represent the mean PSI across replicates. Mouse developmental timepoints are shown on the X-axis, and tissues on the Y-axis. (B) SUZ12 exon 4 PSI quantification in single cells across human and mouse early developmental stages. Each dot represents an individual cell. The number of cells analysed is shown in brackets. The upper, centre and lower lines of the boxplot indicate the 75%, 50%, and 25% quantiles, respectively. Whiskers extend to the most extreme data point within 1.5× the interquartile range. (C), Heatmap of percentage of sequence identity of different vertebrates species at 200 bp upstream of intron 3 plus the first 15 bp of exon 4 (left), exon 4 (centre) and the last 15 bp of exon 4 plus 200 bp downstream of intron 4 (right). Sequence alignments contain 159 sequences from 153 different vertebrate species. (D) Sequence logos of the SUZ12 exon 4 and the six neighbouring intronic nucleotides of eutherian mammals (n = 80 species), marsupials and monotremata (n = 11 species), and all other non-mammalian vertebrates (n = 68 sequences). Grey vertical lines highlight the most variable (i.e. not conserved) exonic nucleotides between constitutively spliced and AS species. Positions on the X-axis start at the first exonic base (uppercase nucleotides). Upstream and downstream intronic sequences (lowercase nucleotides) are numbered with negative and positive numbers, respectively. Information content is expressed in bits on the Y-axis. Species phylogeny and AS patterns are shown on the right.

**Supplementary Figure 2.**
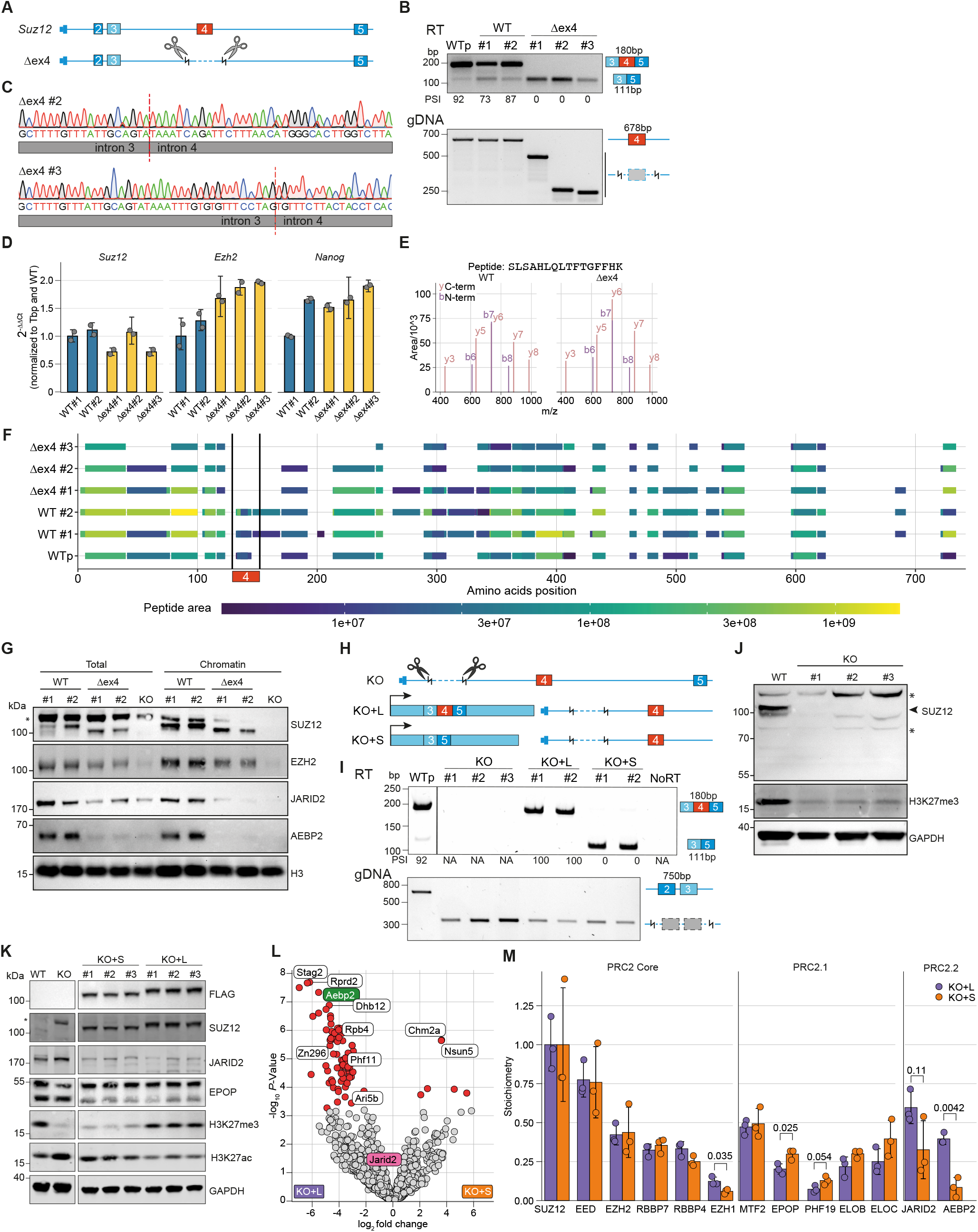
Cell line characterisation and KO+L/S interactome. (A), Schematic representation of CRISPR-Cas9 deletions performed in ESCs to generate Δex4 cell lines. (B), RT-PCR validating Suz12 exon 4 splicing (top); and genomic DNA PCR validating Suz12 exon 4 deletion (bottom). PSI, shown below top gel. The; expected amplicon size is shown on the right. (C) Sanger sequencing chromatogram and intronic sequence flanking Suz12 exon 4 in Δex4 ESC. (D) Bar plot qPCR quantification of Suz12, Ezh2, and Nanog in Δex4 and WT control ESC. (E) Parallel reaction monitoring (PRM)-based targeted mass spectrometry spectra of the Suz12S peptide identified from WT and Δex4 ESCs. (F) SUZ12 IP-MS protein peptide coverage. The location of exon 4 is marked by vertical black lines; the colour bar shows the peptide peak area. (G) Chromatin fractionation WB of WT, Δex4 or KO ESCs. (H) Schematic representation of the CRISPR/Cas9 deletions performed in ESCs to generate the KO, KO+L and KO+S cell lines. (I) RT-PCR validating Suz12 KO, KO+L, and KO+S exon 4 splicing (top); genomic DNA PCR validating Suz12 exon 2-3 deletion (bottom). (J) WB of Suz12 KO ESCs. The arrowhead indicates the correct SUZ12 band. (K) WB of total protein extract from KO+S and KO+L ESCs. (L) Volcano plot of SUZ12 IP-MS of KO+L vs KO+S ESCs. Significant proteins (adj. *P*-value *≤* 0.05, *log*_2_ FC > |1.5|) are shown in red. Data were analysed as in Figure 2d. AEBP2 and JARID2 are coloured as in Figure 2b. (M) Stoichiometric ratio of KO+L and KO+S calculated as in Figure 2E. Points indicate biological replicates (n = 3 clonal cell lines). Error bars represent sd. Statistics: t-test. In G, J and K, asterisks indicate unspecific bands.

**Supplementary Figure 3.**
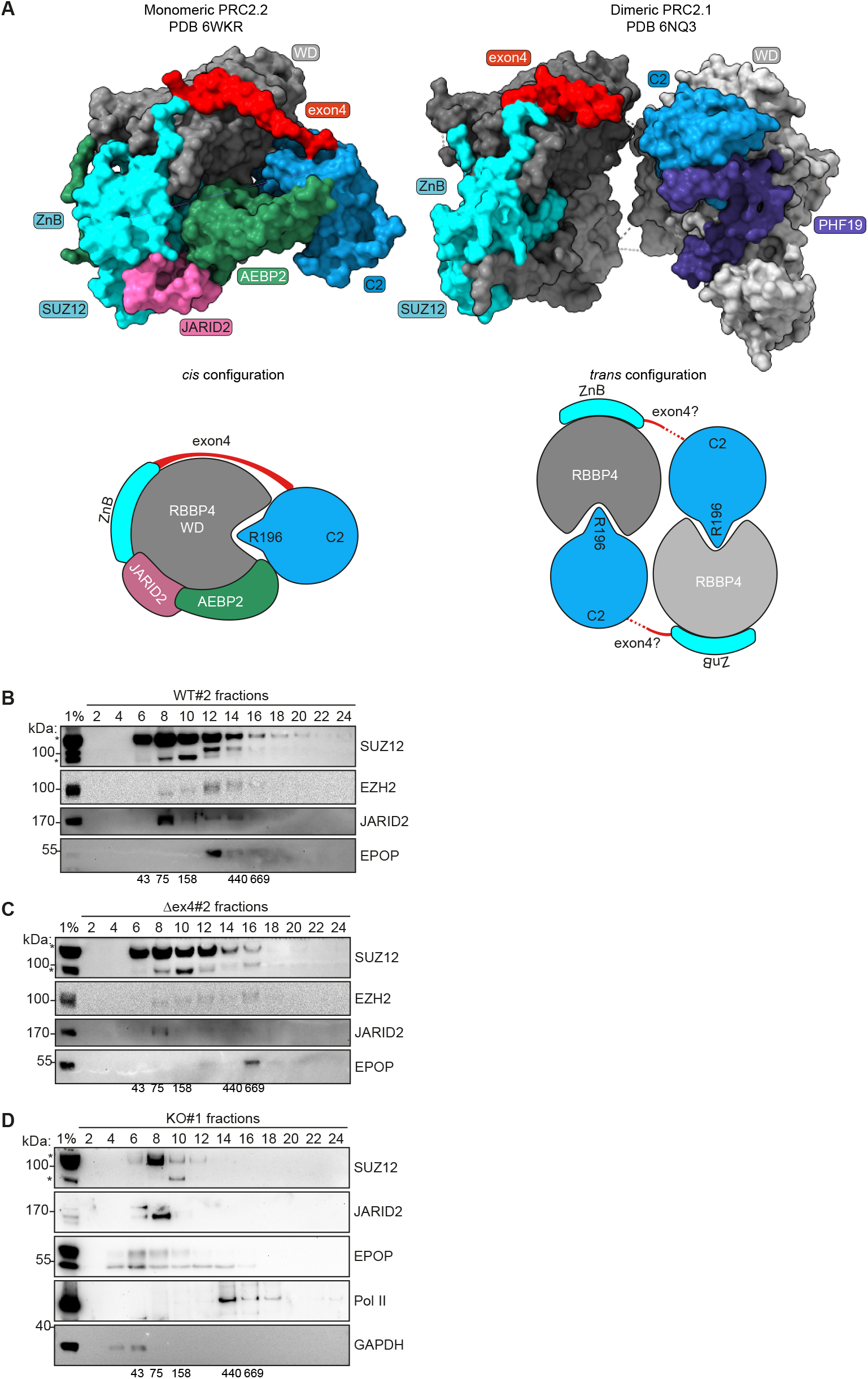
SUZ12-S favours PRC2 dimerization. (A), Top: Surface rendering of monomeric PRC2.2 (left) according to ^7^ and dimeric PRC2.1 (right) according to ^14^ Bottom: schematic representation of cis (left) and trans (right) ZnB-exon4-C2 configuration. Exon 4 dashed line indicates low resolution uncertainty in the structure. (B–D), Glycerol gradient fractionation WB of WT (B), Δex4 (C), and KO cells (D). Higher fractions correspond to higher molecular weights. Numbers below gels correspond to expected fraction molecular weight (kDa). In (B–D), asterisks indicate unspecific bands.

**Supplementary Figure 4.**
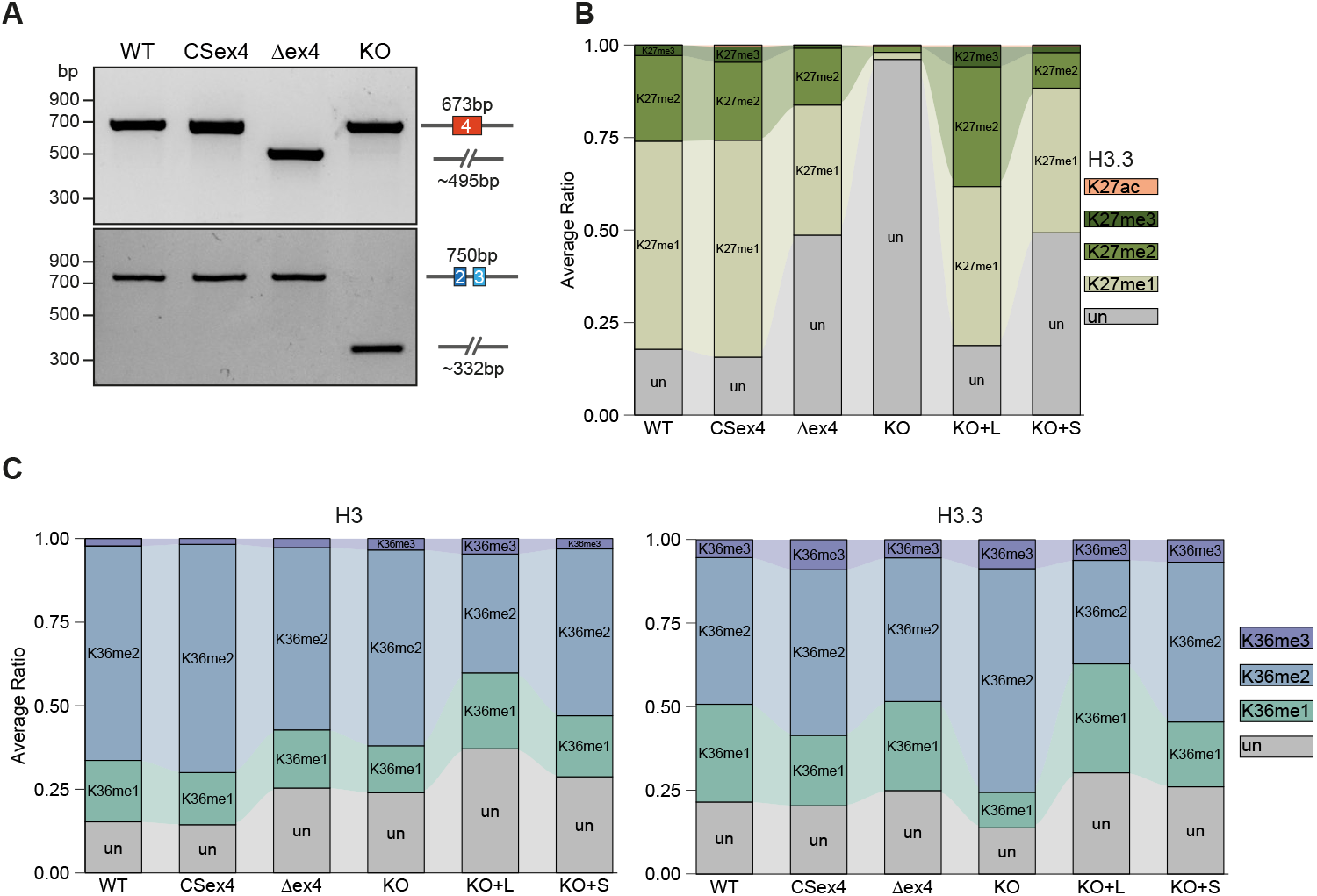
Characterisation of global and local H3K27me3 deposition. (A) Genomic DNA PCR of exon 4 locus (top) and exon 2–3 (bottom) in ESCs. (B,C) Quantitative MS of histones H3 and H3.3 in ESCs. The stacked barplot of the average PTM ratio of H3.3 (27-40) is shown in (C), and of the average PTM ratio of H3 (top) and H3.3 (bottom), in (C) (27-40) (analysis EpiProfile). Combinatorial PTMs are deconvoluted and piled-up (see Methods).

**Supplementary Figure 5.**
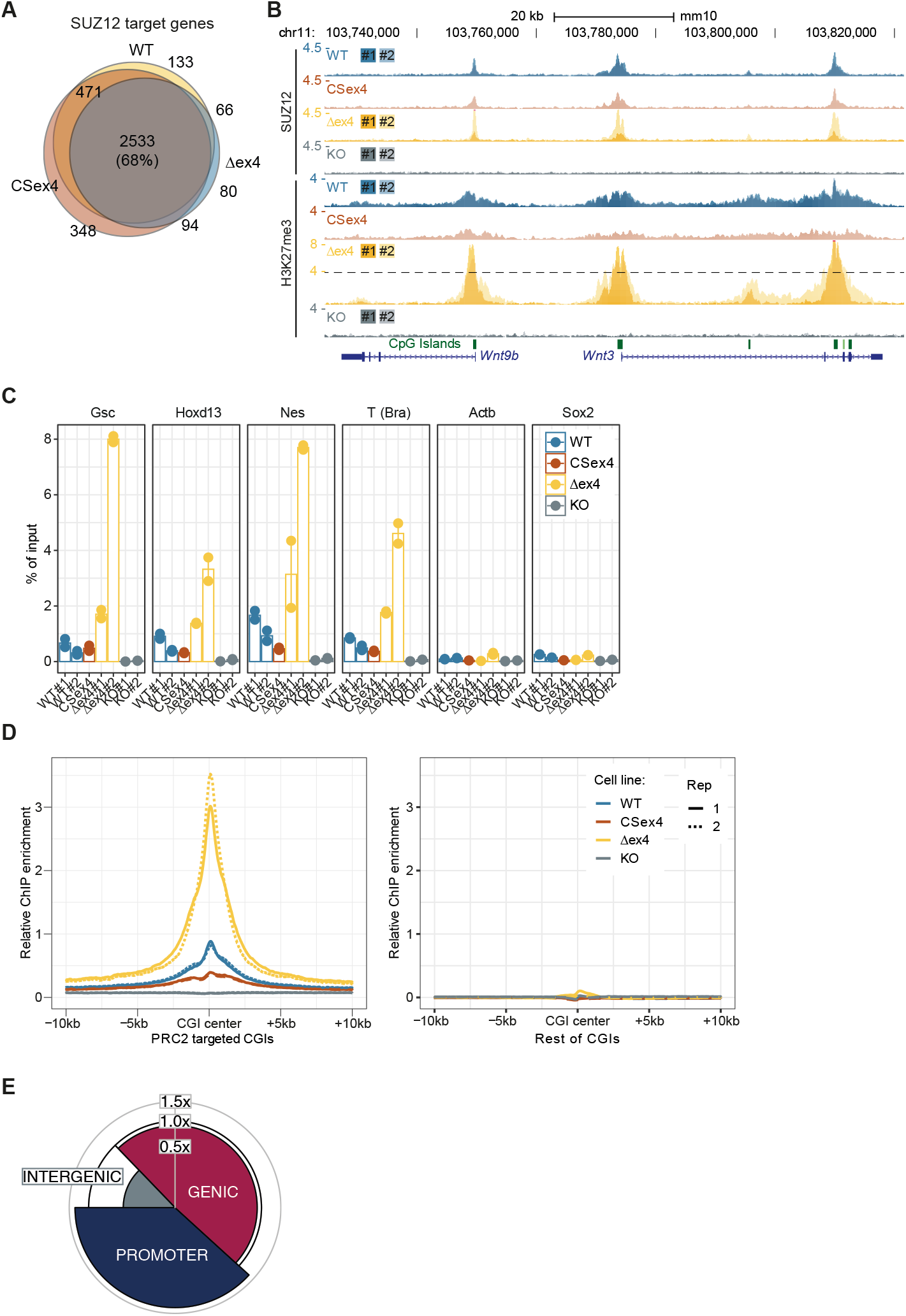
SUZ12 localisation and H3K27me3 deposition upon exon 4 splicing. (A) Venn diagram of Suz12 peaks found in ESCs. (B) ChIP-seq tracks of H3K27me3 in two representative developmental genes targeted by PRC2. CGI location in green. (C) Barplot of ChIP-qPCR enrichment for 4 PRC2 target genes promoters and 2 control genes. (D) Metaplot of library size normalised H3K27me3 at PRC2-targeted CGIs (left) or non-PRC2–targeted CGIs (right). Metaplots are centred on the middle of the CGI (n = 2 clonal cell lines). (E) Spie chart of the percentage distribution of H3K27me3 peaks in Δex4 ESCs. The radial increase corresponds to the fold-change relative to WT ESCs.

**Supplementary Figure 6.**
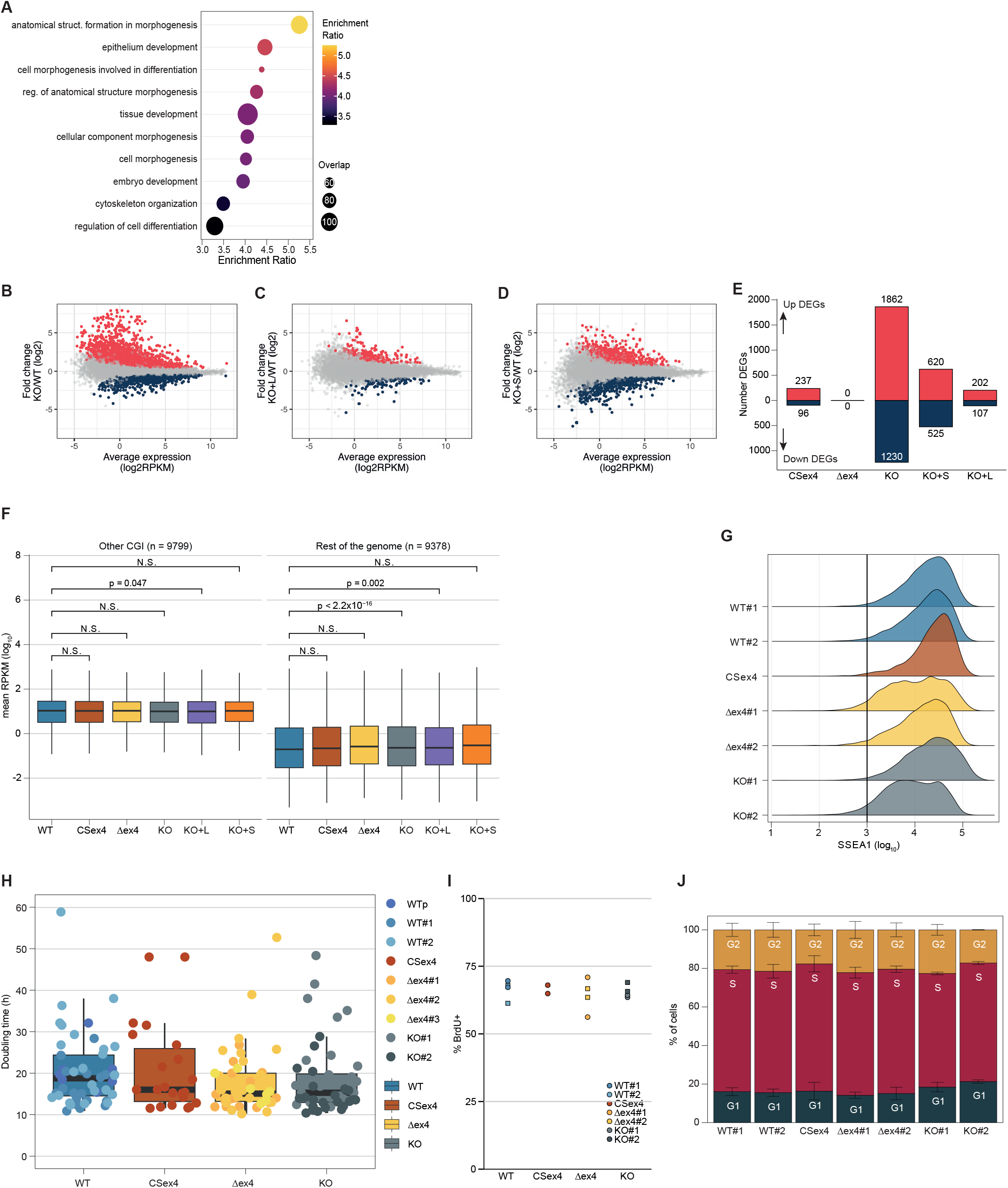
Characterisation of the impact of SUZ12-S and SUZ12-L on gene expression and ESC features. (A) Bubble plot of GO terms statistically enriched in DEGs upregulated in CSex4. (B–D), MA plot of KO (B), KO+L (C), and KO+S (D). (E) Number of up- and downregulated DEGs as compared to WT ESCs. (F) Boxplot of mean expression in ESCs of genes with a CGI not targeted by SUZ12 (n = 9799) and genes without CGIs (n = 9378). Statistics, Wilcoxon test; P-value adjustment, Bonferroni; ns, not significant. (G) Density plot of the signal intensity of SSEA1 staining as measured by FACS. (H) Boxplot of doubling time of ESCs in serum/LIF. (I) Stacked barplot of cell cycle analysis in ESCs, as measured by FACS. (J) Stacked barplot quantifying the percentage of ESCs in each stage of the cell cycle. Error bars: sd.

**Supplementary Figure 7.**
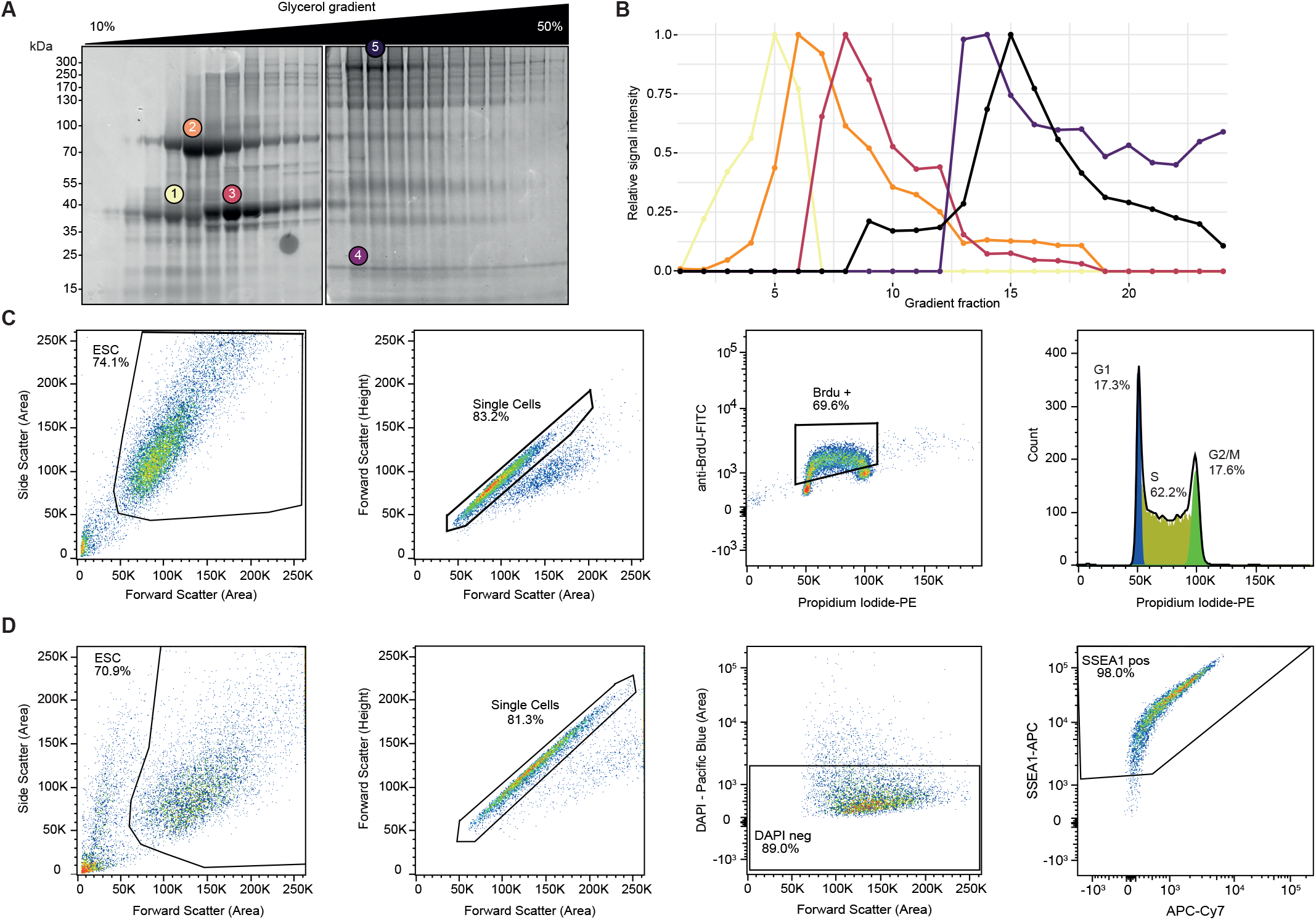
Gradient fractionation calibration and FACS gating strategies. (A,B) Bluesafe staining of the protein complex fractionation weight ladder (A) and its quantification (B). Numbers and colours indicate the following proteins: 1) Ovalbumin (43 kDa, monomer); 2) Conalbumin (75kDa, monomer); 3) Aldolase (40 kDa, tetramer; 4) Ferritin (20 kDa, 24-mer); 5) Thyroglobulin (330 kDa, dimer). (C) Gating strategy for flow cytometry analysis of proliferation rate (BrdU) and Cell cycle profile (propidium iodide). (D) Gating strategy for flow cytometry analysis of SSEA1 surface marker.

## Supplementary Tables

**Supplementary Table 1** | List of publicly available RNA-seq SRA Run IDs used in this study.

**Supplementary Table 2** | List of oligonucleotides sequences used in this study.

**Supplementary Table 3** | Proteomic experiments processed data tables: SUZ12 PRM, IP-MS differential enrichment analysis, Histone MS differential enrichment analysis.

## Word count

~~~
File: Article.tex Encoding: utf8 Sum count: 9097
Words in text: 7934 Words in headers: 85
Words outside text (captions, etc.): 1070 Number of headers: 28
Number of floats/tables/figures: 6 Number of math inlines: 8
Number of math displayed: 0 Subcounts:
    text+headers+captions (#headers/#floats/#inlines/#displayed)
    180+1+0 (1/0/0/0) Abstract
    364+1+415 (1/2/2/0) Section: Introduction
    2058+44+655 (7/4/2/0) Section: Results
    554+1+0 (1/0/0/0) Section: Discussion
    4778+38+0 (18/0/4/0) Section: Methods
~~~

